# Microbiome Analysis of Gut Bacterial Communities of Healthy and Diseased Malaysian Mahseer (*Tor tambroides*)

**DOI:** 10.1101/2021.12.08.471852

**Authors:** Melinda Mei Lin Lau, Cindy Jia Yung Kho, Leonard Whye Kit Lim, Siew Chuiang Sia, Hung Hui Chung, Samuel Lihan, Kasing Apun

## Abstract

**Aims:** The gut microbiota is referred to an ‘extra organ’ and is ciritical in assisting the host in terms of nutrition and immunity. Environmental stressors could alter gut microbial community and cause gut inflammation. This study aimed to investigate and compare the gut microbiota community between healthy and diseased *Tor tambroides*.

**Methodology and results:** In this study, such gut microbial alterations were explored using NGS-based 16S rDNA sequencing on the Malaysian mahseer (*T. tambroides*). Three adult healthy and three diseased adult Malaysian mahseers (showing signs of exophthalmia, coelomic distension and petechial haemorrhage) were obtained from LTT Aquaculture Sdn Bhd. Our results revealed significant differences in microbial diversity, composition and function between both populations of *T. tambroides.* Alpha diversity analysis depicts lower diversity of gut microbiota composition in diseased *T. tambroides* as compared to the healthy group. In particular, *Enterobacteriaceae*, *Aeromonas, Bacteroides, Vibrio* and *Pseudomonas* were found within gut microbiota of the diseased fishes. In addition, cellulose-degrading bacteria and protease-producing bacteria were identified from the gut of *T. tambroides*.

**Conclusion significance and impact of study:** Thus, our findings emphasised on the association between the alteration in gut microbiota composition and infectious abdominal dropsy (IAD) in *T. tambroides.* This finding is important to provide basic information for further diagnosis, prevention and treatment of intestinal diseases in fish.

## INTRODUCTION

Within the past few decades, study of microbiota within gastrointestinal tract (GIT) had been achieving a remarkable progress with the discovery of more GIT microbiome communities on a host organism (Liu *et al.,* 2016; Li *et al.,* 2016; Egerton *et al.,* 2018; Tran *et al.,* 2018; Butt and Volkoff, 2019; Tan *et al.,* 2019; Burtseva *et al.,* 2021). As of today, there are two main approaches to discover on gut microbiota community: culture-dependent microbiological methods and culture-independent methods. The classical method involved seeding gut sample directly on either selective or universal media (Hovda *et al.,* 2007; Tarnecki *et al.,* 2017) while the later involved DNA barcoding. For instance, denaturing gel electrophoresis, qPCR and fluorescence in situ hybridization (Hovda *et al.,* 2007; Tarnecki *et al.,* 2017; Egerton *et al.,* 2018). As culture-dependent methods are time-consuming and selective, it is unable to provide the entire microbial diversity of complex environments. (Hovda *et al.,* 2007). Thus, NGS-based method involving metabarcoding based on 16S rRNA gene is now a popular method among researchers to undercover more uncultured forms of microorganisms and estimate different bacterial groups within the sample as it is able to describe both cultivable and uncultivable bacteria (Tarnecki *et al.,* 2017; Egerton *et al.,* 2018).

Gut microbiota is considered as ‘extra organ’ due to its important role in intestinal development, immunological protection, growth and health and homeostasis (O’ Hara amd Shanahan, 2006). Thus, various studies comparing the gut microbiota composition between healthy and diseased freshwater fish including largemouth bronze gudgeon (*Coreius guichenoti*) suffering from furunculosis (Li *et al.,* 2016), Crucian carp (*Carassius auratus*) suffering from “red-operculum” disease (Li *et al.,* 2017) and grass carp (*Ctenopharyngodon idellus*) suffering from enteritis had been done to further understand on its role. Myriad diversity of mutualistics, commensal and pathogenic microbes within the intestinal tube would assist the host in terms of protection against infectious agents, nutrients uptake and absorption as well as synthesizing digestive enzymes (Nayak, 2010). Researchers concur that the intestine acts as the main portal of entry for pathogen invasion and disease occurrence (Dash *et al.,* 2008; Nayak, 2010; Li *et al.,* 2016). Environmental stress such as poor water quality, exerted upon the fish would disrupt the gut microbiota community and cause gut inflammation (Liu *et al.,* 2016). An inflamed gut can compromise the immune system and eventually cause overgrowth of invading pathogenic bacteria and existing opportunistic pathogens (Zeng *et al.,* 2017). Massive development of *Enterobacteriaceae* initiated by gut inflammation would induce further pathogen invasion as well (Zeng *et al.,* 2017). Intestinal diseases in fishes encompassing infectious dropsy (Dash *et al.,* 2008; Aly and Ismail, 2016), enteritis (Tran *et al.,* 2018), ‘Red-Operculum’ Disease (Li *et al.,* 2017), furunculosis (Li *et al.,* 2016), are reported to be associated with changes in gut microbiota community. Invasion of pathogenic species is associated with imbalance microbiota community. For instance, infectious dropsy (*Aeromonas hydrophilia* and *Pseudomonas fluorescens*) (Aly and Ismail, 2016), enteritis (*Pseudomonas* and *Flavobacterium*) (Tran *et al.,* 2018), ‘Red-Operculum’ disease (*Vibrio, Aeromonas* and *Shewanella*) (Li *et al.,* 2017) and furunculosis (*Aeromonas salmocida*) (Li *et al.,* 2016).

*Tor tambroides* is one of the most exorbitant freshwater species in Malaysia due to its unique flesh contributed by its engkabang consumption. It is still remained re-evaluated as it is classified as data deficient by International Union for Conservation of Nature (Kottelat *et al.,* 2018). Nevertheless, it is under threat due to environmental degradation caused by human activities (Pinder *et al.,* 2019). Among the *Tor* species, *T. tambroides* which is indigenous to Sarawak of east Malaysia, is one of the most valuable freshwater species. To date, there are 16 *Tor* species that can be found worldwide with three (18.8%) found in Malaysia, which are *Tor tambroides, Tor tambra* and *Tor douronensis* (Ng, 2004). Studies had found out that different colours of *T. tambroides* (silver-bronze and reddish) had the possibility to associate with environmental influences (Esa *et al.,* 2006; Siraj *et al.,* 2007). However, ambivalent descriptions of these three *Tor* species inhabiting in Malaysia had cause misidentification among scientific communities in past few years (Walton *et al.,* 2017). Furthermore, a recent study in disentangling the phylogenetic relationship among *T. tambra* and *T. tambroides* sampled from Malaysia (Sarawak, Pahang and Terengganu) and Indonesia (Java) using 13 protein coding genes and two ribosomal RNA genes had shown a monophyletic clustering based on the sampling locations (Lim *et al.,* 2021). *T. tambroides* is among the most valuable game fish, aquaculture fish and ornamental fish which can also serve as a superior source of protein (Ng, 2004), with an aquaculture production of Mahseer in 2018 and 2017 recorded at 12.31 tonnes (RM 3.43 million) and 24.19 tonnes respectively (RM 5.07 million) (DOF, 2018; 2017). Thus, it can be said that *Tor* genus species owns great potential in aquaculture industry (Ingram *et al.,* 2005).

Similar to other freshwater fishes, *T. tambroides* is vulnerable to infectious diseases as well. Infectious abdominal dropsy (IAD) disease is one of the infectious diseases faced by most freshwater fish due to bacterial infection. It is described as an acute haemorrhagic disease which causes mortality and morbidity among fish (Dash *et al.,* 2008). IAD had been reported across various species of fishes, including Indian major carps (*Catla catla, Labeo rohita, Cirrhinus mrigala*), common carp (*Cyprinus carpio*), goldfish (*Carassius auratus*), grass carp (*Ctenopharyngodon idella*), bighead carp (*Aristichthys nobilis*), silver carp (*Hypojtjalmichthys molitrix*), crucian carp (*Cyprinus carassius*), tench (*Tinca tinca*), sheatfish (*Silurus glanis*) and rainbow trout (*Onchorhynchus mykiss*) (Dash *et al.,* 2008; Petty *et al.,* 2012). As one of the most common genera dominating freshwaters, *Aeromonas* also associated with virulence genes and haemolytic activity which allows infection when the host in under stress (Hamid *et al.,* 2016; Vatsos, 2017). Besides mahseers, it was being detected in giant freshwater prawns, tilapia and catfish as well (Chiew *et al.,* 2019).

To the best of our knowledge, there were various studies conducted to understand more on *T. tambroides,* but most of them focused more on its feed formulation and feed additives in improving its growth rate while only a minority of them focused on the molecular biology aspect (Apun *et al.,* 1999; Ishak *et al.,* 2016; Kamarudin *et al.,* 2012; Misieng *et al.,* 2011; Ng *et al.,* 2008; Ng & Andin, 2011; Ramezani-Fard *et al.,* 2012; Lau *et al.,* 2021; Lim *et al.,* 2021). Although there is a finding highlighting on the gut microbiota comparison among wild and captive *T. tambroides* sampled from West Malaysia (Tan *et al.,* 2019), the gut microbial community of *T. tambroides* suffering from disease remains unclear Thus, in this study, Illumina MiSeq Paired-end sequencing (Apical Scientific Sdn Bhd) was used to sequence the V3-V4 regions of 16S rRNA genes for gut bacterial identification on healthy and diseased *T. tambroides* sampled from aquaculture farm at in Sarawak, East Malaysia, in order to identify and compare the gut microbiota between healthy and dropsy diseased *T. tambroides* and to identify the possible pathogenic agents associated with the diseased state of the fish. V3-V4 regions were chosen in this study as it generates a higher richness and diversity while giving a more in-depth characterization of microbial composition (García-López *et al.,* 2020).

## MATERIALS AND METHODS

### Sample Collection

In this study, both healthy and diseased sample groups were included, with three adult fish representing each group. All the samples were obtained from an aquaculture farm located at Asajaya, Sarawak, Malaysia (GPS coordinates: 1° 32’ 53.647’’ N, 110° 32’ 53.233’’ E). Altogether, three adult healthy *T. tambroides* (standard length 44.4 cm ± 0.43 cm, weight 1.05 kg ± 0.15 kg) and three diseased *T. tambroides* (standard length 57.57 cm ± 6.08 cm, weight 2.77 kg ± 0.71 kg) were obtained. The diseased *T. tambroides* fishes were examined morphologically to record on its clinical symptoms before some internal fluids were cultured.

### Fish Dissection and DNA Extraction

Fishes were euthanized and their abdomens were dissected using sterile instruments inside the HEPA filter graded laminar flow hood. All procedures in this study were in compliance to the guidelines and permission approved by the Animal Ethics Committee of Universiti Malaysia Sarawak (UNIMAS/TNC(PI)-04.01/06-09(17)). The gut was taken from oesophagus to anus. The GIT was cut into small pieces of an approximate length of 1 cm within CTAB buffer before DNA extraction using modified CTAB-based protocol (Chung, 2018). Eluted DNA was taken to check for its concentration using a Nanodrop DS-11 Series (DeNovix; USA). Only samples with ratio absorbance at 260 nm and 280 nm (A_260/280_) around 1.8 were selected for further processing.

### Library Preparation and Sequencing

The V3-V4 hypervariable regions of 16S rRNA genes of gut microbiota was chosen to be amplified through Polymerase Chain Reaction (PCR) using following primers: 27F_B (5’ TCGTCGGCAGCGTCAGATGTGTATAAGAGACAG 3’) and 519R_A (5’ GTCTCGTGGGCTCGGAGATGTGTATAAGAGACAG 3’). 27F_B is the forward primer and 519R_A is the reverse primer which were both widely used (Lane, 1991; Handi *et al.,* 2011). The reaction mixtures (25 µL) included: 2X KAPA HiFi HotStart ReadyMIx (12.5 µL) (Kapa Biosystems, USA), forward and reverse primers (1 µM and 5 µL each), and template DNA (5 ng). Amplification conditions were set as follow: 3 min of initial denaturation at 95 °C, followed by 25 cycles of denaturation at 95 °C for 30 s, annealing at 55 °C for 30 s, elongation at 72 °C for 30 s, and a final elongation at 72 °C for 5 mins.

The DNA samples were sequenced on an Illumina MiSeq platform. In brief, DNA samples were subjected to library preparation prior to sequencing. The amplicons were cleaned up for the attachment of unique index adapter pairs to the amplicons using Nextera XT Index kit (Illumina, USA). The indexed DNA libraries were cleaned up with Agencourt AMPure XP (Beckman Coulter, USA). The concentrations of libraries were quantified using Qubit dsDNA HS Assay Kit and Qubit 2.0 Fluorometer (Thermo Fisher Scientific, USA) and size validated using Agilent 2100 Bioanalyzer (Agilent, USA). Next, the libraries were normalised and pooled for subsequent MiSeq sequencing (2 x 300 bp paired-end). All sequences were submitted to NCBI Sequence Read Archive (SRA) under accession number of PRJNA778601 (https://www.ncbi.nlm.nih.gov/sra/PRJNA778601).

### Data Analysis using Quantitative Insights into Microbial Ecology (QIIME)

Data was then analysed using Quantitative Insights into Microbial Ecology (QIIME2 ver 2020.8) (Bolyen *et al.,* 2019). Adapter sequences were cleaved from both paired-ends forward and reverse reads using cutadapt command prior to trimming chimeric sequences. Divisive Amplicon Denoising Algorithm 2 (DADA 2) was used to denoise and filter chimeric sequences based on parametric model which infer true biological sequences from reads (Prodan *et al.,* 2020). Forward and reverse reads were denoised independently and merged prior to removal of chimeric sequences through ‘removeChimeraDenovo’, which eventually formed Amplicon Sequence Variants (ASV). DADA2 was chosen due to its high sensitivity to detect and differentiate ASVs at single-base resolution, even at high abundance ratio (Callahan *et al.,* 2017; Prodan *et al.,* 2020). ASVs-based method had demonstrated high sensitivity as well and specificity as good or even better than Operational Taxonomic Unit (OTU) methods which gave better differentiated ecological patterns (Callahan *et al.,* 2017). Clustered ASVs were then summarized into different taxonomic levels based on GreenGenes database at 99% identity threshold (version 13_8) (DeSantis *et al.,* 2006).

Alpha rarefaction curve was plotted to determine the sequencing depth. Different alpha diversity index parameters (Chao1, Simpson, and Shannon) were utilized to further elucidate on the species richness and diversity of each sample among gut microbiota of both healthy and diseased *T. tambroides.* The value of Chao1 index reflects theoretically predicted richness and abundance (Chao, 1984). Simpson index ranges from 0 to 1 with 0 indicating infinite diversity and 1 indicating zero diversity (Simpson, 1949) while Shannon index reflects species richness and evenness (Shannon, 2001). Furthermore, in order to estimate the coverage of total species represented in each sample, Good’s Coverage was taken into account as well. Beta diversity analysis aims to evaluate diversity among gut microbiota of healthy and diseased *T. tambroides.* Thus, principle coordinate analysis (PCoA) was plotted for visualization of differences or likeliness of gut microbiota diversity among both healthy and diseased *T. tambroides* based on phylogenetic or count-based distance metrics. Weighted UniFrac was chosen in this study for PCoA as it is able to detect the differences in relative abundances of each taxon within the communities (Lozupone *et al.,* 2007). Permutational multivariate analysis of variance (PERMANOVA) was used to evaluate statistical differences in beta diversity while Kruskal-Wallis test was employed to evaluate differences in alpha diversity index of gut microbiota diversity among both healthy and diseased *T. tambroides*.

ANCOM was used to determine the differences of gut microbial communities among both healthy and diseased *T. tambroides* (Mandal et al., 2015). ANCOM was selected over other statistical tests as it made no assumptions and performed well even involve thousands of taxa (Mandal et al., 2015). Mandal et al. (2015) stated that the false discovery rate (FDR) was controlled at a desired nominal level which further improved its performance.

## RESULTS AND DISCUSSION

### DNA quality check

The PCR product was visualised on a 1.7% TAE agarose gel at 100V for 65 min (Figure S1). Discrete bands were observed at size ranges of 450 bp to 550 bp, indicating the presence of V3-V4 region of 16S rRNA gene in all the six samples of *T. tambroides,* with an expected size of approximately 460 bp (Figure S1). Only DNA samples with D(260)/D(280) reading more than 1.8 were subjected to 16S metagenetic sequencing (Table S1).

**Table 1.**
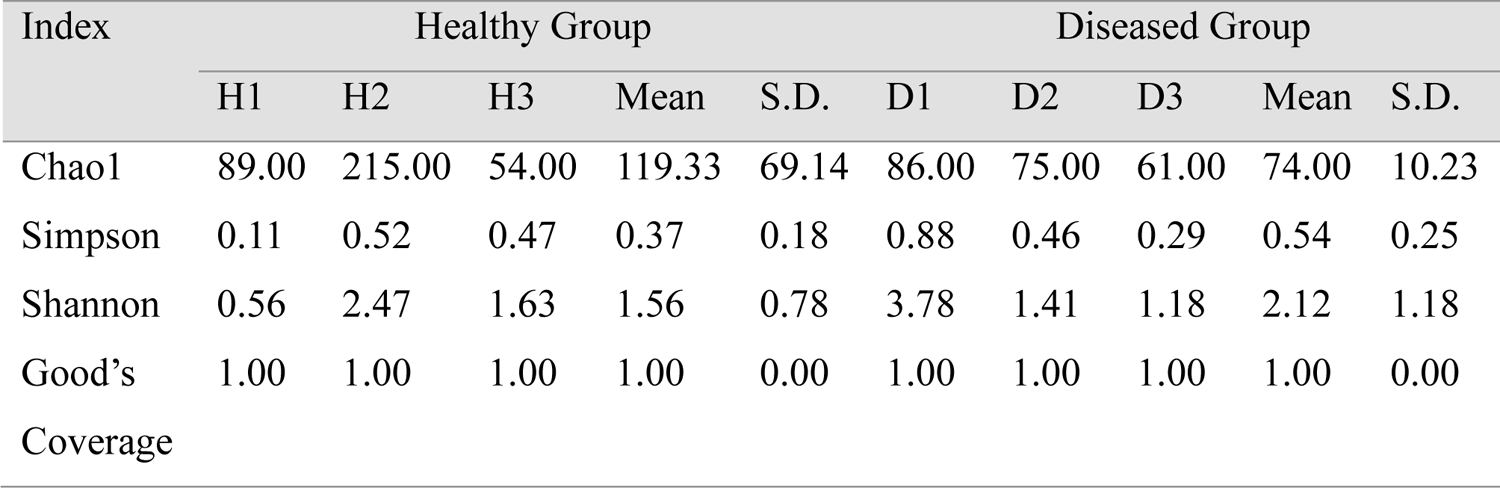
Summary of the alpha diversity of healthy and diseased *T. tambroides* gut microbiota. (S.D. stands for standard deviation)

#### Clinical findings on diseased *T. tambroides*

Diseased adult *T. tambroides* were observed and identified for clinical signs. Some noticeable signs displayed by the three samples include exophthalmia (pop-eye), coelomic distension and petechial haemorrhage on skin, gills, tails and eyes can be observed (Figure 1). Sample D2 showed some loss of scales at its abdominal section as well. Internally, fluid was observed within the internal cavity of diseased *T. tambroides*, which might be due to organ inflammation (Figure 1). A brief microbial culture and 16S sequencing revealed the presence of *Aeromonas hydrophila* in body fluid.

**Figure 1.**
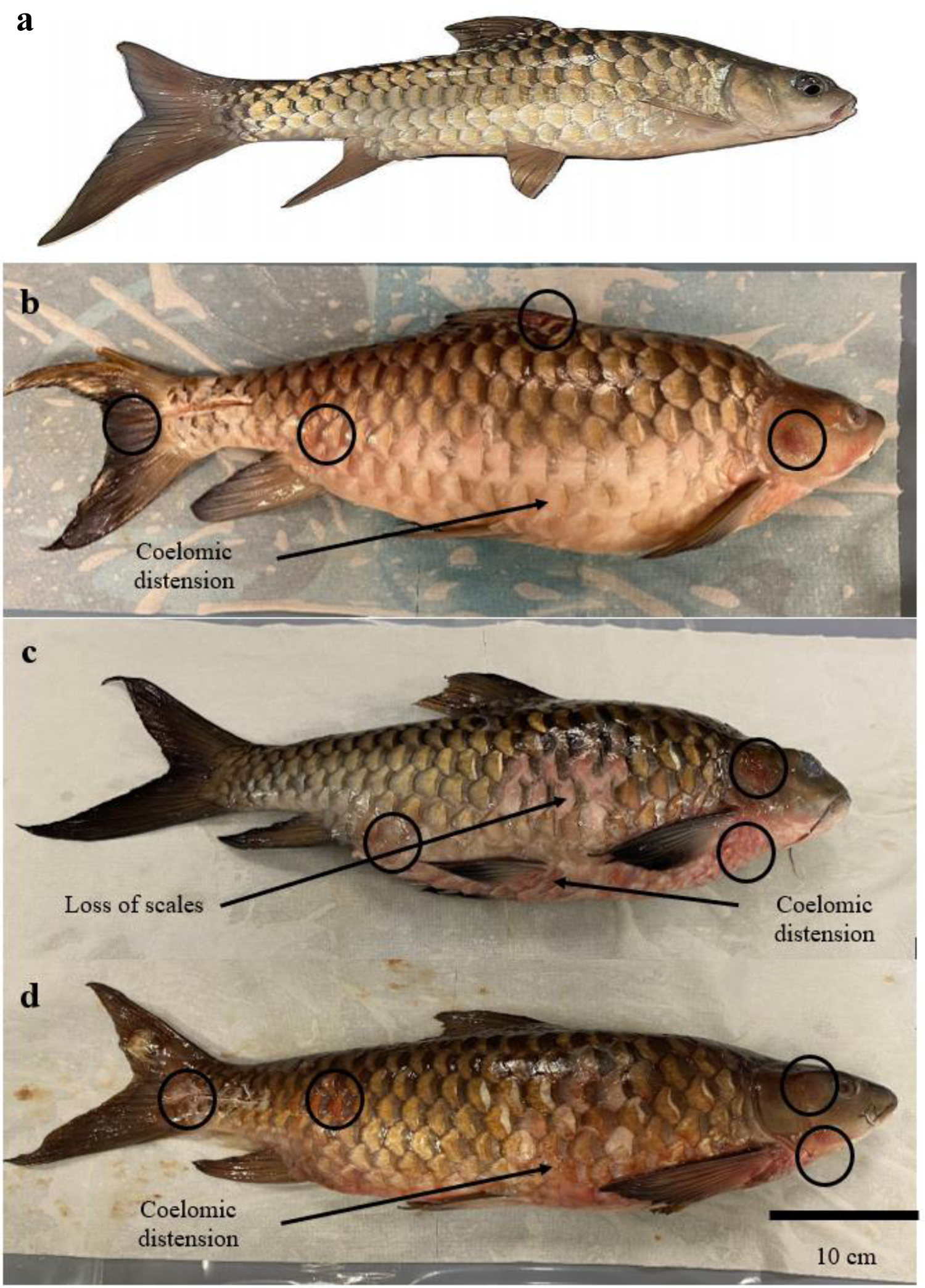
Healthy and diseased samples of *Tor tambroides*, including (a) healthy sample; (b) sample D1 and (c) sample D2; (d) sample D3, with the clinical symptons labelled, including loss of scales and coelomic distension pointed out by arrows while petechial haemorrhage circled on gills, skin, eyes and tails.

In previous studies (Aly and Ismail, 2016; Dash *et al.,* 2008; Lopamudra and Nayak 2020), researchers had reported on the morphological signs of infectious dropsy in *Cyprinus carpio* (common carp), *Catla catla* (South Asian carp), *Labeo rohita* (rohu), *Cirrhinus mrigala* (Mrigal carp) and *Hypopthalmichthys molitrix* (silver carp). The typical clinical symptoms included haemorrhagic lesions presented on skin, fins, tail and eyes, coelomic distension, loss of scales and exophthalmia (Aly and Ismail, 2016). *Aeromonas hydrophilia* and *Pseudomonas fluorescens* were suggested as causative agents for the disease (Aly and Ismail, 2016; Dash *et al.,* 2008).

Studies done on fish disease exhibiting syndrome including exophthalmia, haemorrhagic lesions on fins and abdominal swelling with visceral fluid in Nile tilapia (*Oreochromis niloticus*) had included *Aeromonas veronii, Flavobacterium columnare, Plesiomonas shigeloides, Streptococcus agalacticae* and *Vibrio cholerae* as predominant bacteria (Dong *et al.,* 2015). Clinical signs had been mimicked by infecting the fishes with *A. veronii* and *F. columnare* (Dong *et al.,* 2015). Subsequently, Dong et al. (2017) had identified the bacterial isolates as *Aeromonas jandaei* and *Aeromonas veronii* in diseased Nile tilapia. They revealed a reduced 10- and 100-fold dose of both *Aeromonas* (*Aeromonas jandaeii*: 3.7 x 10^6^ CFU fish^-1^; *Aeromonas veronii*: 8.9 x 10^6^ CFU fish^-1^) had shown diseased symptoms including a significant amount of yellowish fluid built up internally. Dong et al. (2017) emphasised that the similar signs were exhibited with and without the treatment of *Aeromonas* strain.

Our study had revealed genus *Vibrio, Streptococcus, Pseudomonas* and *Aeromonas* in gut microbiota of *T. tambroides* through 16S rRNA gene metagenomic sequencing (Table S6). *Aeromonas* was detected through microbial culture and 16S Sanger sequencing. Nevertheless, *Flavobacterium* was only found in healthy gut microbiota while *Pleisodomonas* was found absent in both groups. As comparison, *Vibrio, Pseudomonas* and *Streptococcus* were found to have greater and non-significant relative abundance in diseased gut microbiota of *T. tambroides. Vibrio* and *Aeromonas* might be the main opportunistic bacteria which are pathogenic to fishes (Li *et al.,* 2017). They were highly-adhesive which enables them to colonise at the intestinal surface mucosa, eventually cause the gut as the primary location for stress-induced infection (Namba *et al.,* 2007). Thus, we suggested *Vibrio* and *Aeromonas* to be associated with the disease-state of *T. tambroides.* However, more evidences are needed to support this postulation. It is suggested that possible pathogens might be present in the intestine prior to disease occurrence and this is in accordance with previous studies done by Li et al. (2016) on largemouth bronze gudgeon suffering from furunculosis. The further invasion of pathogenic strain can be due to various factors, including environmental stressors (pollutants, decreased water quality) which weakens the host’s immune system. This was further supported by a study on microbiota composition analysis of gut in infected Crucian carps and also its surrounding environment by Li et al. (2017). They emphasised on the water physiochemical factors was correlating significantly with the gut microbiota of the fish. Among the water parameters taken, Li et al. (2017) highlighted that the temperature and total ammonia-nitrogen (NO_3_^−^-N) were the most essential factors in shaping the gut microbiota composition.

### Healthy *T. tambroides* owns greater gut microbiota species richness

Illumina Miseq 16S sequencing at the V3-V4 region of 16S rRNA gene enabled an in-depth view and characterization of the gut microbial communities of both healthy and diseased *T. tambroides*. The sequence information of gut microbiome of both healthy and diseased *T. tambroides* were summarised in Table S2. A total of 3,093,610 reads had been generated in this study, ranges from 373,170 to 444,833 and 102,759 to 117,066 in healthy and diseased group respectively, which were found to be higher than previous studies involving fish gut metagenomic studies (Li *et al.,* 2016; Tran *et al.,* 2018; Burtseva *et al.,* 2021). The reads were assigned to 421 ASVs were generated by QIIME 2 at 99% similarity levels, with 301 and 165 ASVs detected in healthy and diseased *T. tambroides* gut microbiota respectively. Among the 301 ASVs detected in healthy *T. tambroides* gut, 12 ASVs (4%) were found to be present in all healthy samples while 15 ASVs (9.10%) out of total 165 ASVs can be found across all diseased samples. In this study, ASV was generated despite of OTU which can be found in other metagenomic studies. Due to the generation of exact sequence by distinguishing the sequence variants differing by one nucleotide, the total number of ASVs generated were lower than OTU which is also dependent on the denoising approach used (Callahan *et al.,* 2017; Nearing *et al.,* 2018).

### Reduced microbiota diversity in dropsy diseased *T. tambroides* gut

The sequencing depth was normalized to 50,000 for both healthy and diseased *T. tambroides* samples (Figure 2) as plateau was observed when the curve flattened gradually as the sequencing depth increase, indicating the current sequencing depth is sufficient to reflect diversity in each sample and the probability of discovering more ASVs beyond the depth is negligible. To estimate completeness of bacterial diversity by obtained 16S rRNA amplicon data, it is necessary to analyze the relationship between Chao1 index and the observed taxa abundance. In Table 1, Good’s coverage had confirmed the sequencing had covered up to an approximate 100% of all gut microbiota found in both healthy and diseased *T. tambroides.* Other alpha diversity parameters (Chao1, Shannon, and Simpson) had been included to evaluate on species diversity of each sample group. Shannon index showed that diseased *T. tambroides* gut microbiota had higher species richness than that of the healthy *T. tambroides.* Nevertheless, Chao1 index and Simpson index for healthy *T. tambroides* which recorded as 119.33 and 0.37 respectively, indicates a higher diversity than diseased fish at 74.00 and 0.54 respectively.

**Figure 2.**
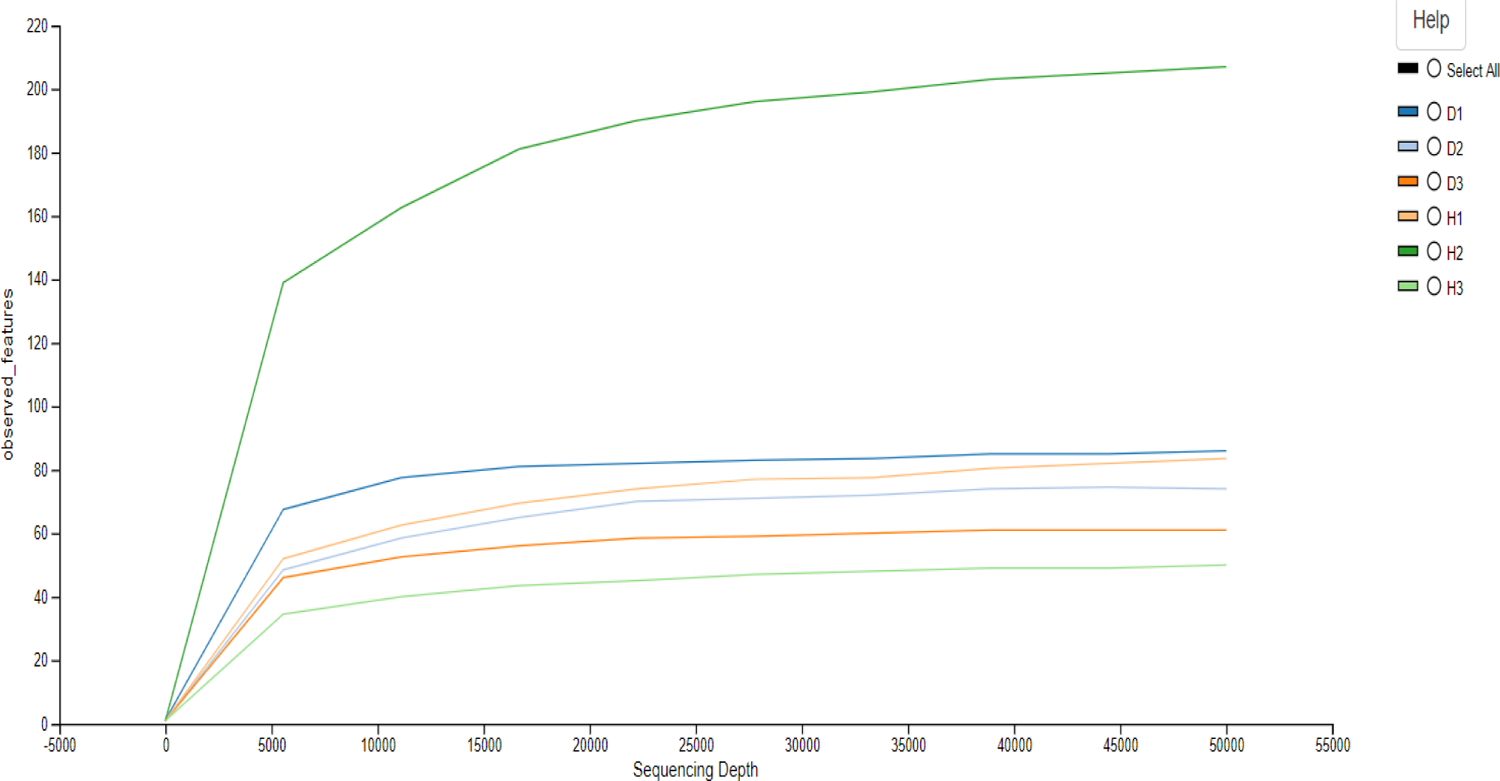
The alpha rarefaction curves of healthy and diseased *T. tambroides* gut microbiota. The x-axis shows the sequencing depth and the y-axis shows the observed features. (H1-H3: biological replicates of healthy *T. tambroides*; D1-D3: biological replicates of diseased *T. tambroides*)

It is in agreement with the diversity resistance hypothesis that greater diversity microbial community possess greater possibility of having a species with antagonistic trait towards invading pathogens (Fargione and Tilman, 2005). Similar trend can be seen in previous studies done on Crucian carp (*Carassius auratus*) (Li *et al.,* 2017), largemouth bronze gudgeon (*Coreius guichenoti*) (Li *et al.,* 2016), Gibel carp (*Carassius gibelio*) (She *et al.,* 2017) and ayu (*Plecoglossus altivelis*) (Nie *et al.,* 2017), exerting greater protection on the host towards invading pathogen. Possible explanation on this is due to competition among invading pathogens with gut commensals, therefore reducing diversity in diseased fish (Xiong *et al.,* 2019). Thus, it can be said that lower gut microbiome diversity is closely associated with diseases (Li *et al.,* 2017; Xiong *et al.,* 2019). However, our findings contradict with previous study on grass carps with intestinal disease (Tran *et al.,* 2018), which might be due to differences in species and associated diseases.

### Beta diversity analysis of healthy and diseased *T. tambroides*

The PCoA plots shown in Figure 3 portrayed clusters based on healthy and diseased samples observed at Principal Coordinate 1 vs Principal Coordinate 2 (PC1 vs PC2). Permutational multivariate analysis of variance (PERMANOVA) had shown non-significant differences among both groups of samples (p = 0.395). However, from Figure 3, the plots of the diseased samples were dispersed widely while the healthy sample plots tend to be located in closer proximity. The overall distribution distance between gut microbiota diversity of healthy and diseased *T. tambroides* was relatively far, further emphasised on the differences among the microbial compositions.

**Figure 3.**
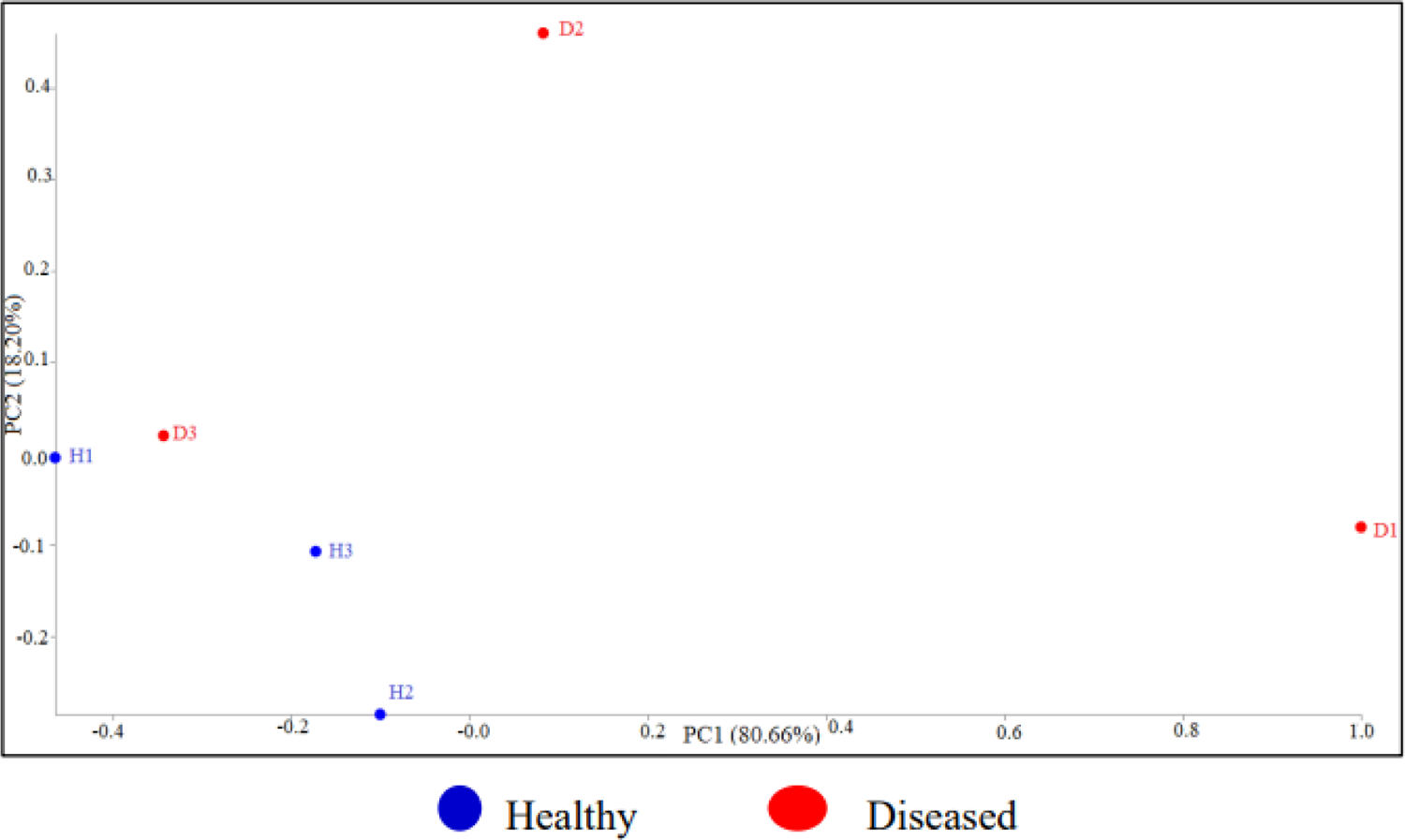
Principal Coordinate Analysis (PCoA) plots based on weighted UniFrac distance metric, representing PC1 vs PC2 on biological replicates of *T. tambroides*

This finding was found to be consistent with reports in grass carps (Tran *et al.,* 2018) suffering from intestinal diseases. Stress exerted upon fishes would disrupt gut microbial community, termed gut dysbiosis, and this is associated with various disorders including inflammatory bowel disease (IBD) and infection (Kamada *et al.,* 2013). Such shift in relative bacterial abundance can be contributed by antibiotic administration, poor water quality and dietary changes. Inflammation would be induced due to compromised immune system, which fosters ‘bloom’ of low-abundance and harmful bacteria. Among the diseased samples, sample D3 tend to lie closer to healthy group samples (Figure 3) than other two diseased samples, suggesting on the less severity in its disease state as compared to D1 and D2.

### Microbiota taxonomy of healthy and diseased *T. tambroides* (Phylum Level)

The gut microbiota of healthy *T. tambroides* was dominated mainly by Proteobacteria (87.397%), followed by Fusobacteria (5.511%), Bacteroidetes (2.844%), Firmicutes (2.235%), unclassified phyla from kingdom bacteria (1.297%) and others (0.715%) (Figure 4). On the other hand, gut microbiota of diseased *T. tambroides* was found to be mainly dominated by Proteobacteria (78.331%), Bacteroidetes (11.470%), Fusobacteria (7.900%), Firmicutes (1.892%) and others (0.412%) (Figure 4). Table S3 had summarized on relative abundance and frequency of each phylum between all samples while Table S4 summarized on the mean and percentage on each phylum on both groups. Proteobacteria was found to be the most abundant phyla in gut microbiota of both datasets. It was due to Proteobacteria are predominant in fish gut across all known metagenomic studies (Tan *et al.,* 2019; Burtseva *et al.,* 2021). The most abundant phylum observed in both datasets were in consistent with the findings by Tan et al. (2019) on wild and captive bred *T. tambroides* sampled from Kenyir Lake.

**Figure 4.**
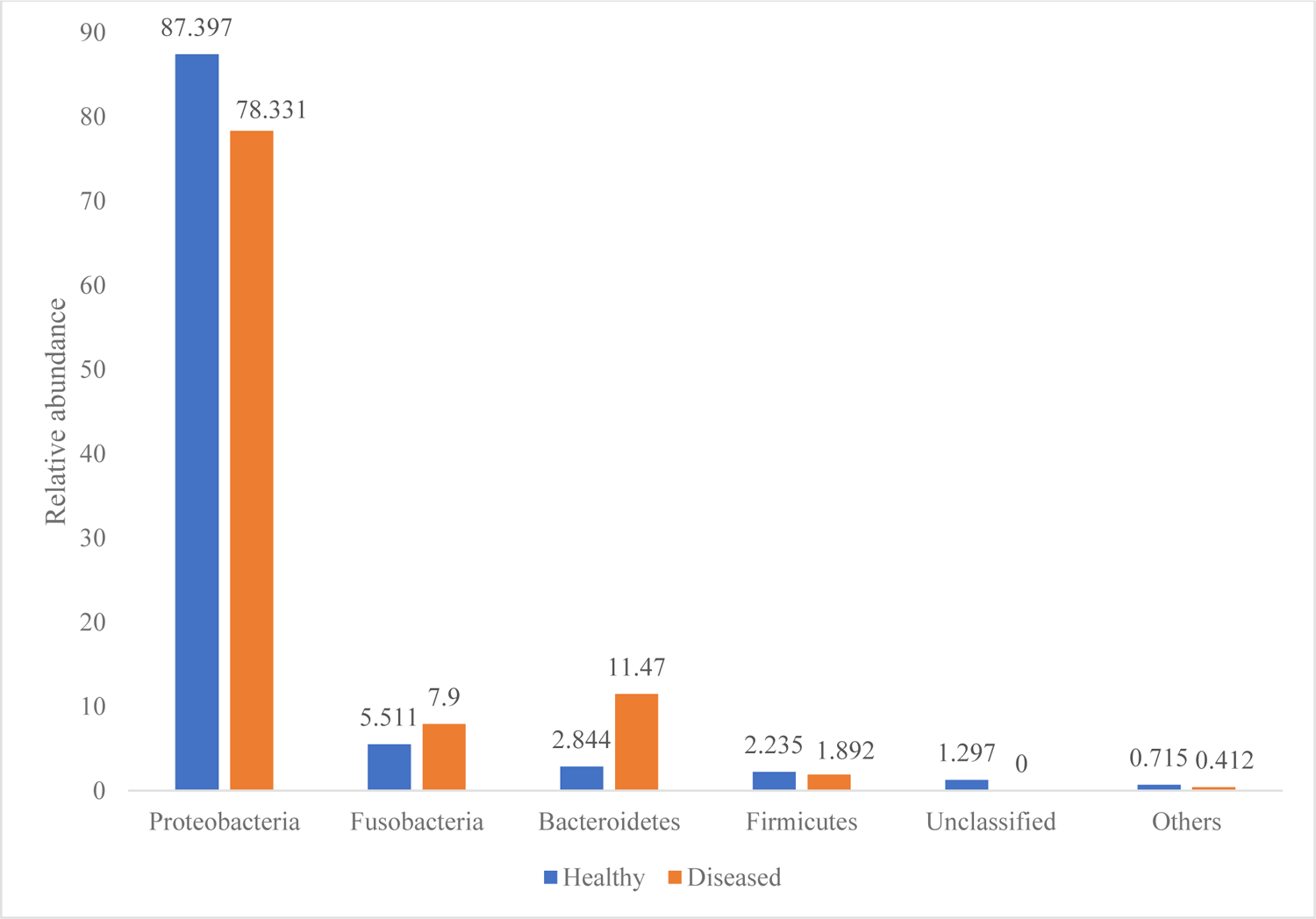
Relative abundance of dominant phyla (%) (mean relative abundance > 0.4%) in gut microbiota of healthy group and diseased group of *T. tambroides.* Phyla with low abundance (< 0.4 %) were combined and categorised under ‘others’.

This study had highlighted an increase in relative abundance of phylum Proteobacteria, Firmicutes and unclassified bacteria which was found in relative higher abundance in healthy gut of *T. tambroides.* In contrast, phylum Fusobacteria and Bacteroidetes in diseased gut microbiota of *T. tambroides* were observed to be in greater relative abundance. Such changes especially with regards to the increased level of the bacterial species within the phyla, suggested that the overgrowth of opportunistic bacteria might reduce the relative abundance of other taxa colonising fish intestine prior to pathogen invasion (Tran *et al.,* 2018). Furthermore, reduction in relative abundance of members within phyla in diseased fish might be related to dietary nutrients accessibility in the fish intestine during diseased state (Tran *et al.,* 2018).

However, the distribution of main classes (Alphaprotebacteria, Betaproteobacteria, Gammaproteobacteria) of phylum Proteobacteria differed among both healthy and diseased group. Alphaproteobacteria (91.77%) had the most counts in healthy gut microbiota of *T. tambroides,* followed by Gammaproteobacteria (8.17%) and lastly Betaproteobacteria (0.06%). In contrast, the dominant class found in diseased gut microbiota is Gammaproteobacteria which accounted for 52.15% of the counts, followed by Alphaproteobacteria (46.93%) and Betaproteobacteria (0.93%). Such phenomenon is due to Gammaproteobacteria consist of a wider range of pathogens as compared to Alphaproteobacteria which associates with host metabolism (Austin & Austin, 2012). Gammaproteobacteria in diseased *T. tambroides* was mostly comprises of possible disease causative agents, such as *Aeromonas, Enterobacteriaceae, Vibrio, Pseudomonas* and *Clostridium,* which is responsible for the diseased state of the fish. ANCOM was performed to determine significant differences among gut microbiota of healthy and diseased *T. tambroides* at phylum level, however there is no significant differences detected. Unclassified bacteria had been identified in a high relative abundance in healthy gut microbiota of *T. tambroides* as compared to the diseased group. It is in consistence with the previous study done on largemouth bronze gudgeon (*Coreius guichenoti*) suffering from furunculosis (Li *et al.,* 2017). In this study, an interesting taxon had been identified, CK-1C4-19, which can be found in healthy *T. tambroides* only, was classified under phylum Tenericutes. It had been reported by previous studies on zebrafish (*Danio rerio*) (Roeselers *et al.,* 2011), fathead minnow (*Pimephales promelas*) (Narrowe *et al.,* 2015) and anjak (*Schizocypris altidorsalis*) (Ghanbari *et al.,* 2019) as small relative abundance.

### Microbiota taxonomy of healthy and diseased *T. tambroides* (Family Level)

There were 72 families identified from both diseased and healthy gut of *T. tambroides* with frequency and relative abundance reported for each sample in Table S5. Out of the total of 72 families, 61 and 50 families were reported within healthy and diseased group respectively. Table 2 showed the relative abundance of top ten most abundant family found in gut of both healthy and diseased *T. tambroides.* Sphingomonadaceae was found to be the dominant family for both datasets, covering up to 79.021% and 36.689% in healthy and diseased group respectively. In previous studies, *Sphingomonadaceae* had been observed in healthy human gut and milk as one of the core members of Proteobacteria and elucidated on its role in host immunity (D’Auria *et al.,* 2013; Corona-Cervantes *et al.,* 2020). *Sphingomonadaceae* was found to be modulating and maintaining immune response (D’Auria *et al.,* 2013) and as a potent stimulators of natural killer T cells (Long *et al.,* 2007). It further stimulates innate immune system to control growth of microbial and as a continuous surveillance of invading pathogens (Long *et al.,* 2007).

**Table 2.**
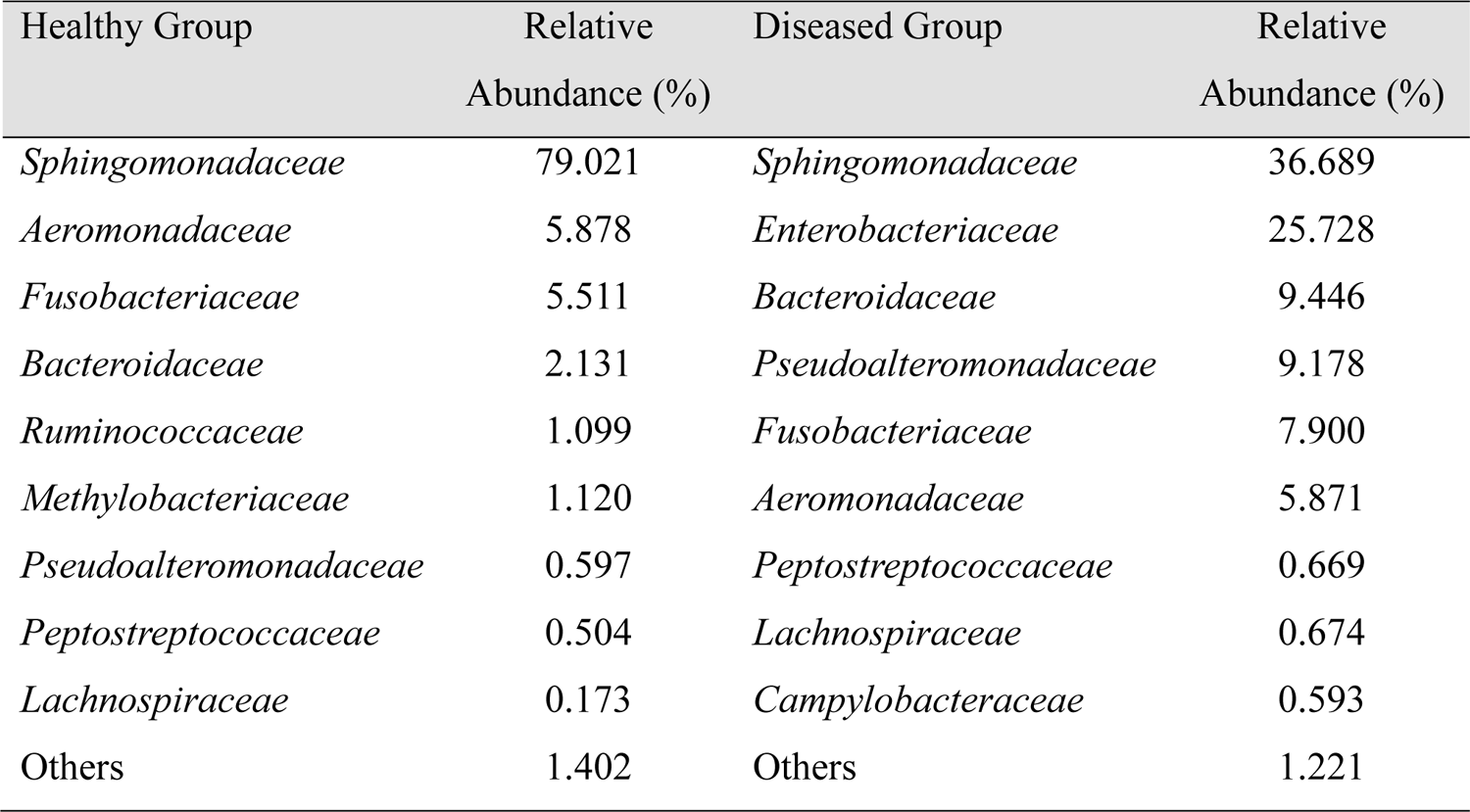
Top ten families found in gut microbiota of healthy and diseased *Tor tambroides* (not including families with lower abundance < 0.40%).

| Healthy Group | Relative<br>Abundance (%) | Diseased Group | Relative<br>Abundance (%) |
| --- | --- | --- | --- |
| *Sphingomonadaceae* | 79.021 | *Sphingomonadaceae* | 36.689 |
| *Aeromonadaceae* | 5.878 | *Enterobacteriaceae* | 25.728 |
| *Fusobacteriaceae* | 5.511 | *Bacteroidaceae* | 9.446 |
| *Bacteroidaceae* | 2.131 | *Pseudoalteromonadaceae* | 9.178 |
| *Ruminococcaceae* | 1.099 | *Fusobacteriaceae* | 7.900 |
| *Methylobacteriaceae* | 1.120 | *Aeromonadaceae* | 5.871 |
| *Pseudoalteromonadaceae* | 0.597 | *Peptostreptococcaceae* | 0.669 |
| *Peptostreptococcaceae* | 0.504 | *Lachnospiraceae* | 0.674 |
| *Lachnospiraceae* | 0.173 | *Campylobacteraceae* | 0.593 |
| Others | 1.402 | Others | 1.221 |

*Enterobacteriaceae* (25.728%) were found to be in high abundance, followed by *Bacteroidaceae* (9.446%) in diseased gut group. In contrast, *Aeromondaceae* (5.878 %) and *Fusobacteriaceae* (5.511%) were found in high abundance in healthy gut of *T. tambroides. Enterobacteriaceae* are the most common overgrown symbionts and can be found in inflammation related infection, such as IBD, obesity and antibiotic treatment (Zeng *et al.,* 2017). This alteration in microbe community compromises colonization resistance which lead to further colonisation and growth of pathogens. High relative abundance of *Enterobacteriaceae* found in diseased gut of *T. tambroides* as compared to the healthy group can be further elucidated through elevation of gut luminal oxygen level due to increased blood flow. Under inflammatory condition, invading microbes were destroyed by transmigrated neutrophils (PMN) accumulating at apical epithelium and gut lumen due to deposition of autoantibodies. NAPDH oxidase were activated to destroy invading pathogens further leads to excessive release of oxygen (Chin and Parkos, 2006). Moreover, both epithelial cells and transmigrated PMNs would produce nitrate species as well. Nitrate respiration favour *Enterobacteriaceae* bloom at certain nitrate-rich tissue environment, as gene encode for nitrate reductase were found within the family. Amount of nitrate would rise significantly when a superoxide reacts with nitric oxide (NO) and eventually converted into nitrate (NO_3_^−^) that enable nitrate respiration (Winter *et al.,* 2013). Such PMN transmigration were shown in an inflammed zebrafish model with expression of green fluorescence protein (GFP) by neutrophils under control of myeloperoxidase promoter (Mathias *et al.,* 2006).

In addition, previous studies had emphasised on bloom of *Enterobacteriaceae* which disrupt gut microbiota balance, lead to gut inflammation and further foster disease occurrence (Zeng *et al.,* 2017). Enterobacterial blooms which involved gram-negative facultative bacteria within class *Gammaproteobacteria* would compromise host immune system, which increases its vulnerability towards incoming pathogens. This can be elucidated through the lipopolysaccharides which act as potent inflammatory pathogen-associated molecular pattern (PAMP) in *Enterobacteriaceae* which worsen intestinal injury (Wallace *et al.,* 2011). *Enterobacteriaceae* bloom act as a key factor for potential pathogenesis development of various diseases. For instance, it would cause the host to be more vulnerable to colitis disease induced by *Clostridium* in mice (Buffie *et al.,* 2012). This is consistent with the increase in relative abundance of *Clostridium* found in diseased gut microbiota of *T. tambroides* as compared to healthy gut.

### Microbiota taxonomy of healthy and diseased *T. tambroides* (Genera Level)

Table S6 had reported on the relative abundance of genera on each samples of both groups. A total of 88 genera was found with 70 found within healthy group and 49 found within diseased group. Both groups shared 30 genera, with 40 and 18 genera uniquely identified from gut microbiota of healthy and diseased *T. tambroides* respectively (Table S7). Figure 5 showed the top ten most abundant genera observed in gut microbiota community of both healthy and diseased *T. tambroides. Aeromonas* (Healthy: 5.878%; Diseased: 5.871%) and *Bacteroides* (Healthy: 2.131%; Diseased: 9.446%) were the top 3 highest abundance in healthy and diseased gut of *T. tambroides. Bacteroides* were found to be increase in abundance in diseased gut as compared to the healthy gut but *Aeromonas* reported on an approximate similar relative abundance in both groups. Furthermore, *Vibrio* (Healthy: 0.600%; Diseased: 9.178%) is reported to be increased in diseased gut microbiota of *T. tambroides,* suggesting it might be associated with the disease-state of the fish. Table 3 showed the top ten most abundant unique observed bacteria among the gut microbiota of both groups. Among them, *Arcobacter* showed the highest relative abundance (0.593%) in diseased gut of *T. tambroides* but it was absent in healthy fish gut.

**Figure 5.**
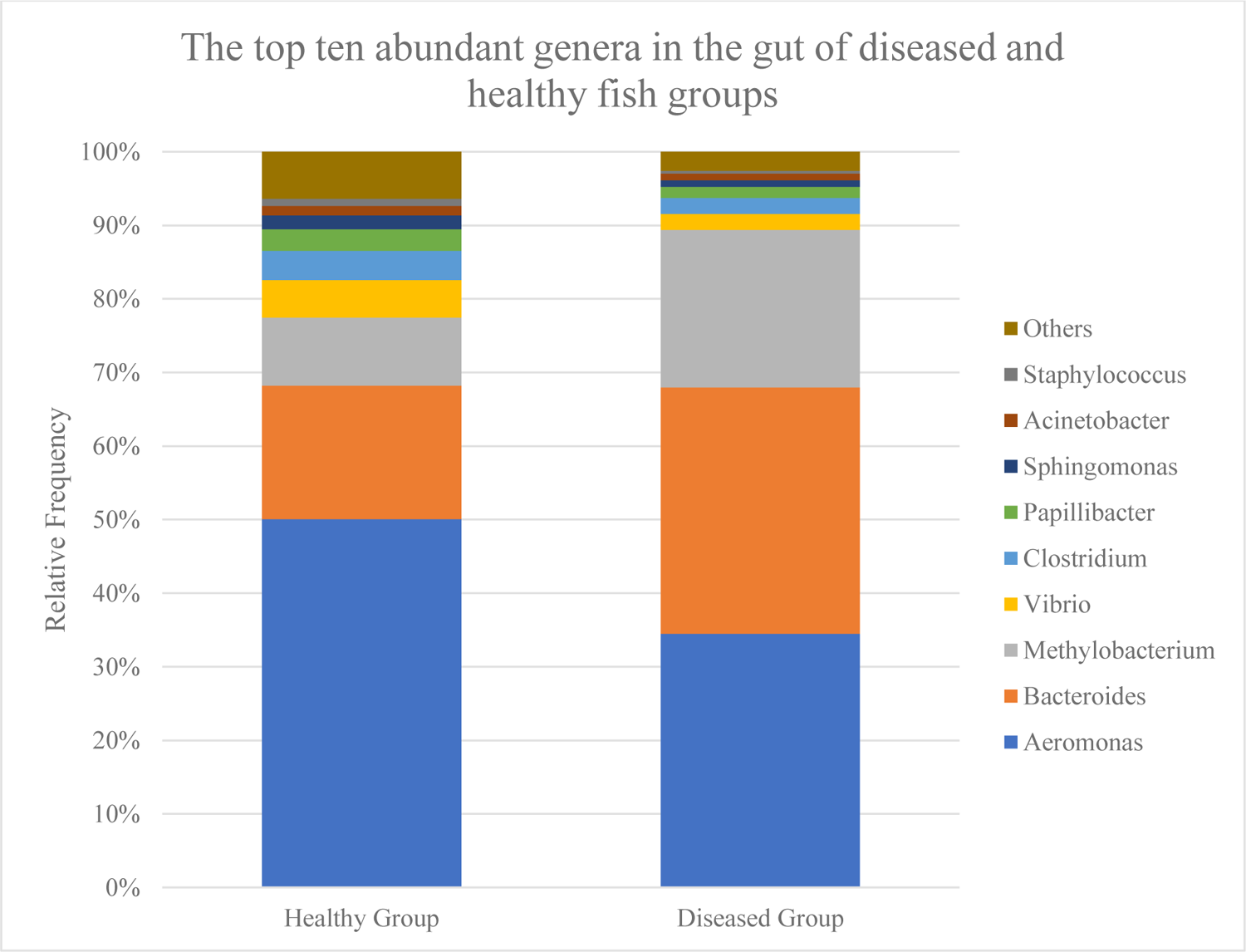
Histogram showing the relative abundance of the top 10 most abundant genera in gu t microbiota of both healthy and diseased groups of *Tor tambroides.* Genera with lower abund ance (< 0.40%) were combined and clustered under ‘others’.

**Table 3.**
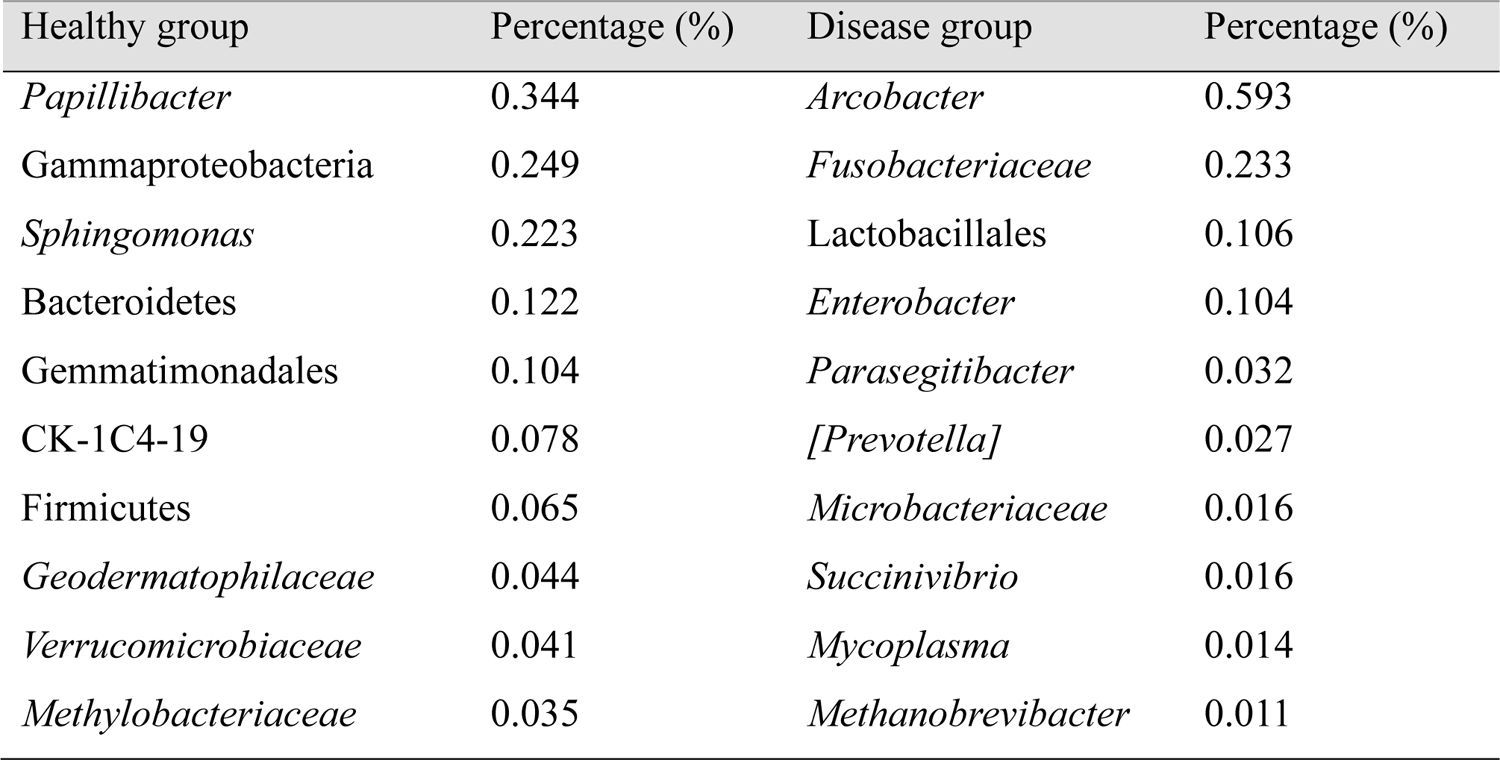
Top ten most abundant unique observed bacteria found in gut microbiota of either healthy or diseased *Tor tambroides*.

In this study, a few highlighted genera have contributed towards the relative abundance in diseased gut as compared to healthy gut of *T. tambroides*, namely *Bacteroides, Vibrio, Pseudomonas* and *Clostridium.* Thus, it can be suggested that these genera may play an essential role in disease progression of *T. tambroides.* Nevertheless, the association between the high relative abundance of these genera and intestinal inflammation remained unclear until further investigation is done (Tran *et al.,* 2018). For instance, an experimental infection with these genera on healthy fish to undercover its association with intestinal inflammation. With the presence of these species in healthy gut as well, it is widely deemed that the fish digestive tract is a reservoir for opportunistic pathogens (Wu *et al.,* 2012).

Probiotics successfully detected in guts of both healthy and diseased *T. tambroides,* include *Streptococcus, Lactococcus,* and from order *Lactobacillales,* which they can be utilized as candidate probiotics. Another possible pathogenic pathogen, *Aeromonas* spp. was found in gut microbiota of both healthy and diseased *T. tambroides,* with a similar relative abundance (Healthy: 5.878%; Diseased: 5.871%). Previous studies had revealed that *Lactococcus* could suppress the microbial activity of seven *Aeromonas* strains (Hagi and Hoshino, 2009). The higher relative abundance of the lactic acid bacteria (LAB) in diseased *T. tambroides* may suggest the presence of higher suppression activity by *Aeromonas* spp. Nevertheless, there were studies suggesting *Aeromonas* play essential role in fish biology rather than as an opportunistic pathogen (Wu *et al.,* 2012). Further *Aeromonas* colonization may be mostly due to stress-induced infection (Wu *et al.,* 2012). Aside from *Lactococcus,* relative abundance of *Streptococcus* is relatively higher in diseased *T. tambroides* and this is consistent with the findings by Tran et al. (2018). It might be involved in controlling the intestinal inflammation in *T. tambroides.* The underlying mechanisms need further investigations.

Elevated production and release of mucin is a hallmark of intestinal inflammation mostly triggered by enteric infection, which act to expulse pathogens and maintain the integrity of mucus layer as the first line of host defence (Frank *et al.,* 2007). Significant increase of *Bacteroides,* as a mucin-degrading bacteria, was detected in diseased gut microbiota of *T. tambroides,* which mediated release of less complex sugar from mucin, such as lactose, raffinose, melibiose and galactinol. *Bacteroides* arbitrated release of sialic acid from mucin in inflamed gut which can be taken up by bacteria for sugar production, further enhance *Enterobacteriaceae* bloom (Stecher, 2015).

*Arcobacter* was found uniquely in gut microbiota of diseased *T. tambroides* as compared to the healthy group. *Arcobacter* which previously known as ‘Aerotolerant *Campylobacter”* was found as a zoonotic pathogen infecting both animals and human (Ho *et al.,* 2006). It is usually associated with gastrointestinal diseases, with high degree of similarity of clinical signs as compared to the *T. tambroides* in this study. The clinical signs include inflammatory gut, fallen scales and rotted fins (Açik *et al.,* 2016). *Arcobacter* had been isolated from intestinal tracts from zebrafish and human, yet its pathogenicity mechanisms were still in vague (Karadas *et al.,* 2013; Açik *et al.,* 2016). Studies had shown on its ability to adhere onto the wall of gut and invade as other bacteria such as Campylobacters (Açik *et al.,* 2016).

Cellulose-degrading bacteria including *Actinomycetospora, Clostridium, Ruminococcus* and *Enterobacter* were found within gut of *T. tambroides,* which is consistent with the finding by Wu et al. (2012). Possible protease-producing bacteria found were *Flavobacterium,* and *Gammaproteobacteria* (Skrodenytė-Arbačiauskienė, 2000). *Flavobacterium* was identified from healthy guts of *T. tambroides* in lower relative abundance but was absent in the diseased group. It was suggested to be associated with various digestive processes by producing enzymes such as amylase, chitinase and protease (Skrodenytė-Arbačiauskienė, 2000).

### Analysis of composition of microbiomes (ANCOM)

Analysis of composition of microbiomes (ANCOM) was used to determine the differences of gut microbial communities among both healthy and diseased *T. tambroides* (Mandal *et al.,* 2015) (Table S8). Both healthy and diseased gut microbiota of *T. tambroides* showed the highest abundance from family *Sphingomonadaceae* (Figure 6), but there are no significant differences shown in both sample groups. As comparison of gut microbiota of healthy group, some observed bacteria experienced a significant increment in their relative abundance of gut microbiota in diseased *T. tambroides,* including *Fusobacteriaceae*, *Bacteroides*, *Vibrio*, Bacteroidales and *Clostridium*. These changes found among gut microbiota suggested that occurrence of diseases would cause changes in gut microbiota composition and diversity. In Figure 6, among the top 15 abundant observed bacteria, only *Enterobacteriaceae* showed significant differences in gut microbiota between diseased and healthy *T. tambroides*.

**Figure 6.**
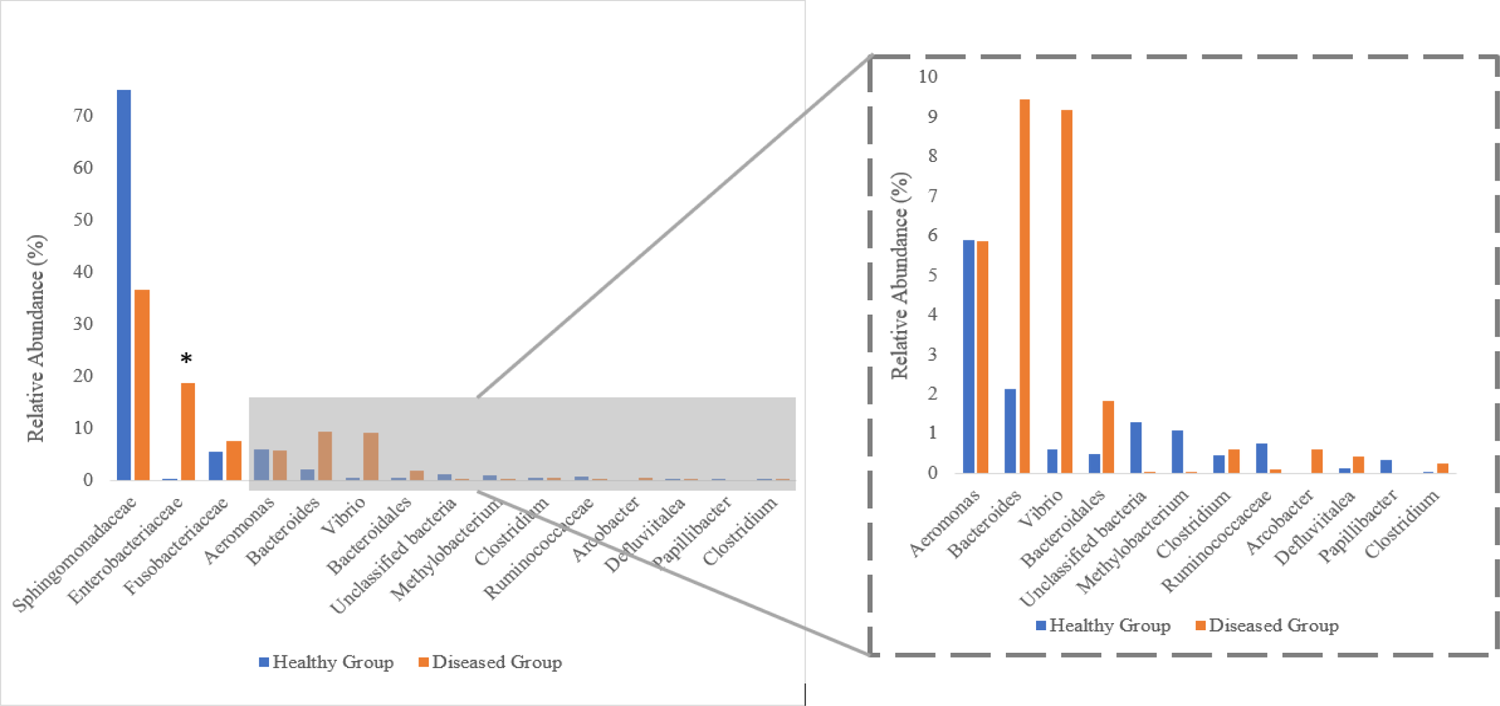
Comparison of top 15 most abundant observed bacteria in gut microbiota of both healthy and diseased *T. tambroides* (Bars with * indicated significant difference between healthy and diseased *T. tambroides* samples through ANCOM.

## CONCLUSION

To our knowledge, this is the first report that delineated the changes of composition, diversity and function of gut microbiota in Malaysian mahseer (*T. tambroides*) suffering from diseases. We compared the healthy and diseased *T. tambroides* gut microbiota and revealed more on the diversity of microbial communities. Based on the beta diversity analysis plotted on PCoA, gut microbiota of healthy *T. tambroides* were generally distantly spaced from the diseased *T. tambroides.* Furthermore, our findings unearthed exclusive genera namely *Vibrio, Bacteroides, Enterobacteriaceae, Pseudomonas, Fusobacteriaceae* and *Clostridium* in diseased gut microbiota of *T. tambroides.* Metagenetic sequencing had revealed on the gut microbiota of both healthy and diseased fish from an aquaculture farm, highlighting its role in further diagnosis, prevention, and developing an ideal approach for better health management of Malaysian mahseer. Besides, we had also provided an insight into etiology of IAD and help to control strategies in intensive aquaculture practice and early detection. Yet, further studies are needed to elucidate more on the gut microbiota existing within captive and wild *T. tambroides* to further understand on the physiological functions, such as growth and disease resistance. Studies on reciprocal interactions between intestinal microbiota and host-produced enzymes in response to microbial alterations are recommended for future studies.

## ACKNOWLEDGEMENTS

This work was fully funded by Sarawak Research and Development Council (SRDC) through the Research Initiation Grant Scheme with grant number GL/F07/SRDC/03/2020 awarded to H. H. Chung.

## AUTHORS’ CONTRIBUTION

Chung, Lihan and Apun conceived of the study conception and experimental design. Lau, Kho and Sia were responsible for acquiring data while Lau, Leonard and Kho analysed and interpreted the data. Lau, Leonard and Kho drafted the manuscript. All authors discussed the results and contributed to the final manuscript.

## SUPPORTING INFORMATION

**Figure S1.**
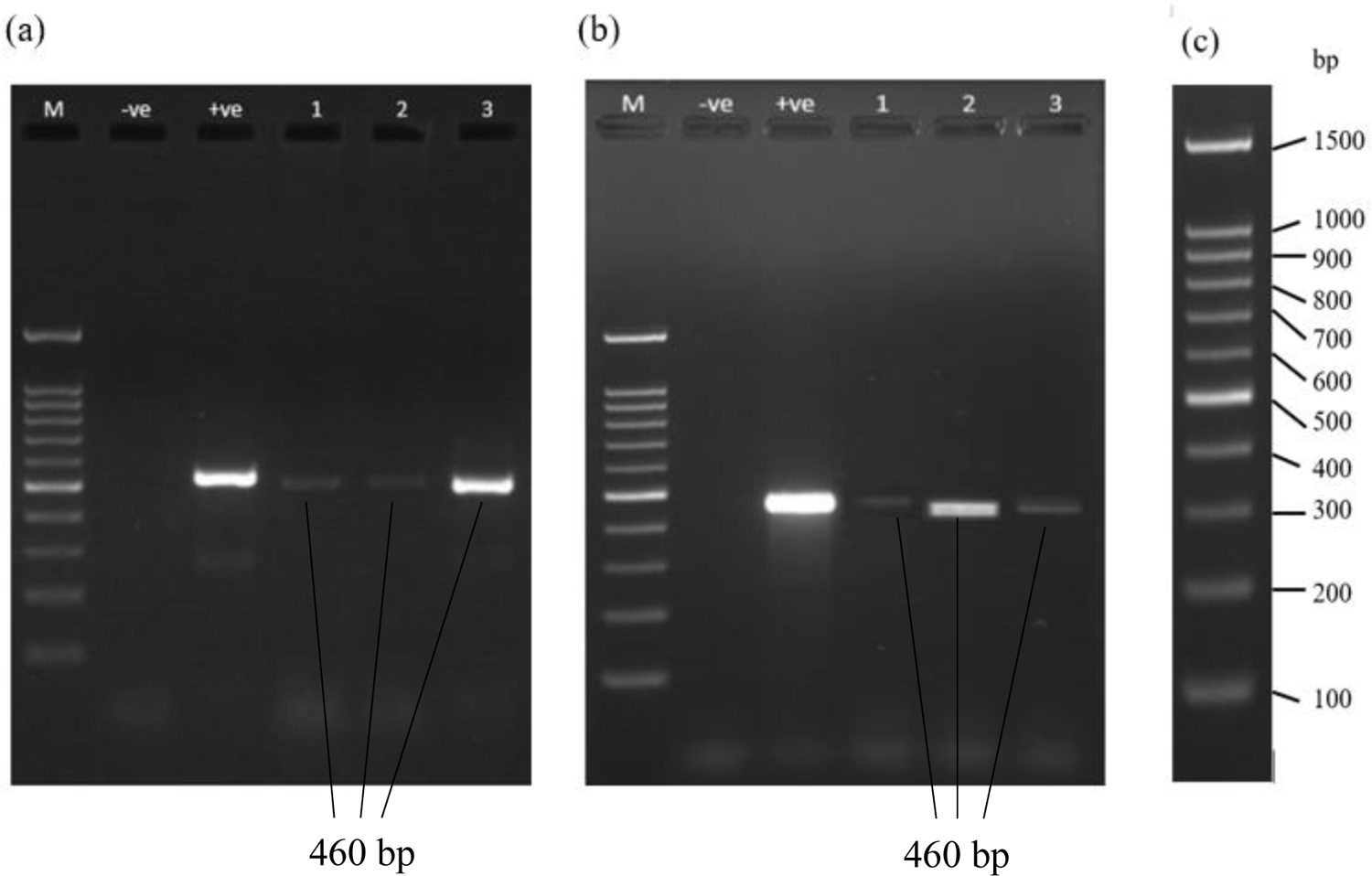
Visualization of PCR products (Forward primer: 5’ TCGTCGGCAGCGTCAGATGTGTATAAGAGACAG and Reverse Primer: 5’ GTCTCGTGGGCTCGGAGATGTGTATAAGAGACAG 3’) under UV on 1.7% TAE agarose gel after running on 100V for 65 min. Lane M is the DNA marker, “-ve” is the no-template control, “+ve” is the positive control with 10 ng of gDNA template for both Figure 1a and 1b. Figure 1a showing the PCR result of healthy fish: Lane 1: H1; Lane 2; H2, Lane 3; H3. Figure 1b showing the PCR result of diseased fish: Lane 1; D1, Lane 2; D2, Lane 3; D3 Figure 1c is the DNA marker used is ExactMark 100bp (C/No. BIO-5130).

---

**Table S1.** DNA Quantification using Nanodrop DS-11 Series (DeNovix; USA)

| Sample | DNA concentration (ng<br>$\mu\text{L}^{-1}$) | Volume ($\mu\text{L}$) | Total amount<br>($\mu\text{g}$) | D(260)/D(280) |
| --- | --- | --- | --- | --- |
| H1 | 117.80 | 13 | 1.53 | 2.002 |
| H2 | 69.25 | 25 | 1.73 | 1.887 |
| H3 | 3564.00 | 20 | 71.28 | 2.123 |
| D1 | 2237.20 | 25 | 55.93 | 2.098 |
| D2 | 786.50 | 20 | 15.73 | 2.043 |
| D3 | 2326.30 | 25 | 58.16 | 2.100 |

H1-H3 stand for the gut samples of healthy fish; while D1-D3 stand for the gut samples of diseased fish.

**Table S2.** Summary of gut microbiome in healthy and diseased *T. tambroides*

| Sample |  | Sequences after<br>merging forward and<br>reverse read | Sequences after<br>QC filter | Sequences after<br>chimera filter | ASVs |
| --- | --- | --- | --- | --- | --- |
| Healthy<br>group | H1 | 398,247 | 277,904 | 274,546 | 89 |
|  | H2 | 373,170 | 257,861 | 250,598 | 215 |
|  | H3 | 444,833 | 304,346 | 277,385 | 54 |
| Diseased<br>group | D1 | 117,066 | 87,079 | 79,097 | 86 |
|  | D2 | 102,759 | 78,046 | 72,957 | 75 |
|  | D3 | 110,730 | 84,549 | 78,366 | 61 |

**Table S3.** Frequency and relative abundance of each phylum for each of the sample, with H1, H2 and H3 stands for healthy *T. tambroides* while D1, D2 and D3 stands for diseased *T. tambroides*.

| Phylum | H1 | % | H2 | % | H3 | % | D1 | % | D2 | % | D3 | % |
| --- | --- | --- | --- | --- | --- | --- | --- | --- | --- | --- | --- | --- |
| Proteobacteria | 269262 | 98.075 | 189278 | 75.531 | 242850 | 87.55 | 33434 | 42.27 | 69777 | 95.641 | 77280 | 98.614 |
| Fusobacteria | 224 | 0.082 | 14620 | 5.834 | 29384 | 10.593 | 17304 | 21.877 | 623 | 0.854 | 276 | 0.352 |
| Bacteroidetes | 502 | 0.183 | 22143 | 8.836 | 179 | 0.065 | 26088 | 32.982 | 163 | 0.223 | 167 | 0.213 |
| Firmicutes | 947 | 0.345 | 16859 | 6.728 | 134 | 0.048 | 1825 | 2.307 | 2107 | 2.888 | 427 | 0.545 |
| Unclassified | 403 | 0.147 | 5461 | 2.179 | 4544 | 1.638 | 25 | 0.032 | 17 | 0.023 | 13 | 0.017 |
| Actinobacteria | 2046 | 0.745 | 499 | 0.199 | 174 | 0.063 | 320 | 0.405 | 208 | 0.285 | 189 | 0.241 |
| Gemmatimonadetes | 833 | 0.303 | 0 | 0 | 0 | 0 | 0 | 0 | 0 | 0 | 0 | 0 |
| Planctomycetes | 133 | 0.048 | 575 | 0.229 | 0 | 0 | 0 | 0 | 36 | 0.049 | 0 | 0 |
| Tenericutes | 0 | 0 | 544 | 0.217 | 85 | 0.031 | 32 | 0.04 | 0 | 0 | 0 | 0 |
| Cyanobacteria | 183 | 0.067 | 161 | 0.064 | 35 | 0.013 | 0 | 0 | 3 | 0.004 | 14 | 0.018 |
| Verrucomicrobia | 0 | 0 | 327 | 0.13 | 0 | 0 | 0 | 0 | 0 | 0 | 0 | 0 |
| Lentisphaerae | 0 | 0 | 120 | 0.048 | 0 | 0 | 0 | 0 | 0 | 0 | 0 | 0 |
| TM7 | 13 | 0.005 | 5 | 0.002 | 0 | 0 | 0 | 0 | 0 | 0 | 0 | 0 |
| Chloroflexi | 0 | 0 | 6 | 0.002 | 0 | 0 | 0 | 0 | 0 | 0 | 0 | 0 |
| Chlamydiae | 0 | 0 | 0 | 0 | 0 | 0 | 27 | 0.034 | 13 | 0.018 | 0 | 0 |
| Euryarchaeota | 0 | 0 | 0 | 0 | 0 | 0 | 24 | 0.023 | 2 | 0 | 0 | 0 |
| Spirochaetes | 0 | 0 | 0 | 0 | 0 | 0 | 18 | 0.023 | 0 | 0 | 0 | 0 |
| OD1 | 0 | 0 | 0 | 0 | 0 | 0 | 0 | 0 | 4 | 0.005 | 0 | 0 |
| Acidobacteria | 0 | 0 | 0 | 0 | 0 | 0 | 0 | 0 | 4 | 0.005 | 0 | 0 |

**Table S4.**
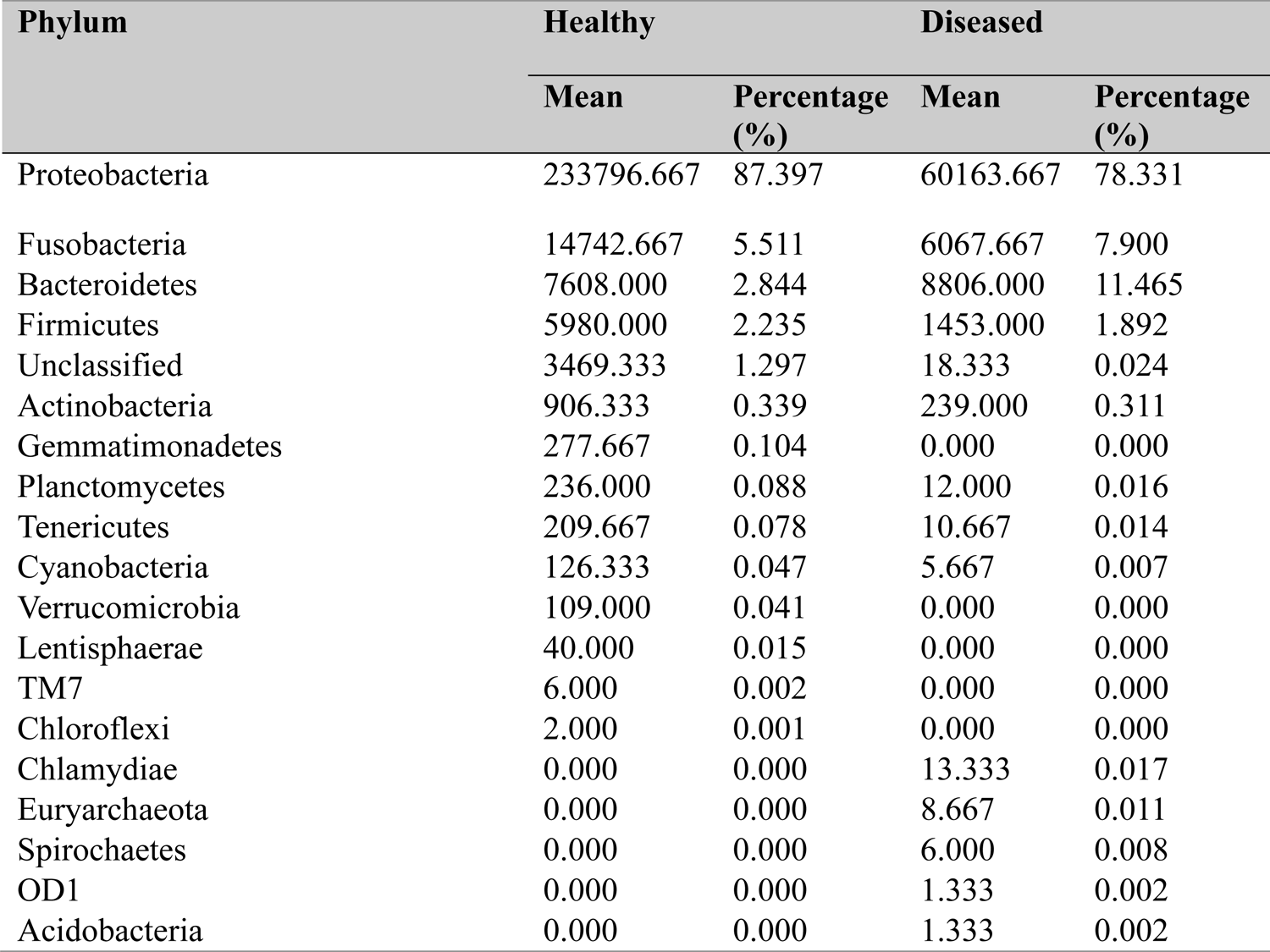
Mean and its percentage for each phylum detected in both healthy and diseased *T. tambroides*.

**Table S5.**
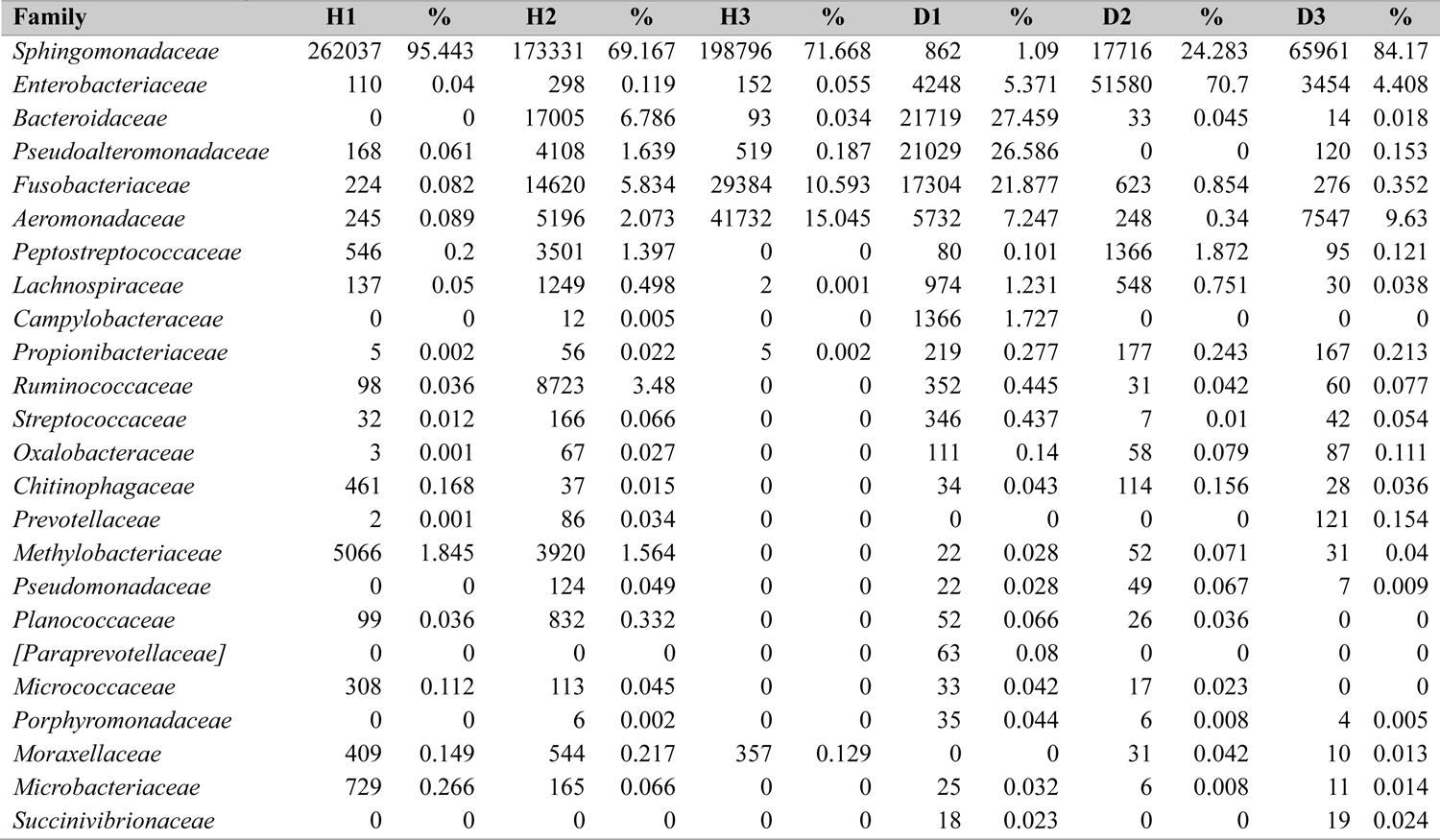
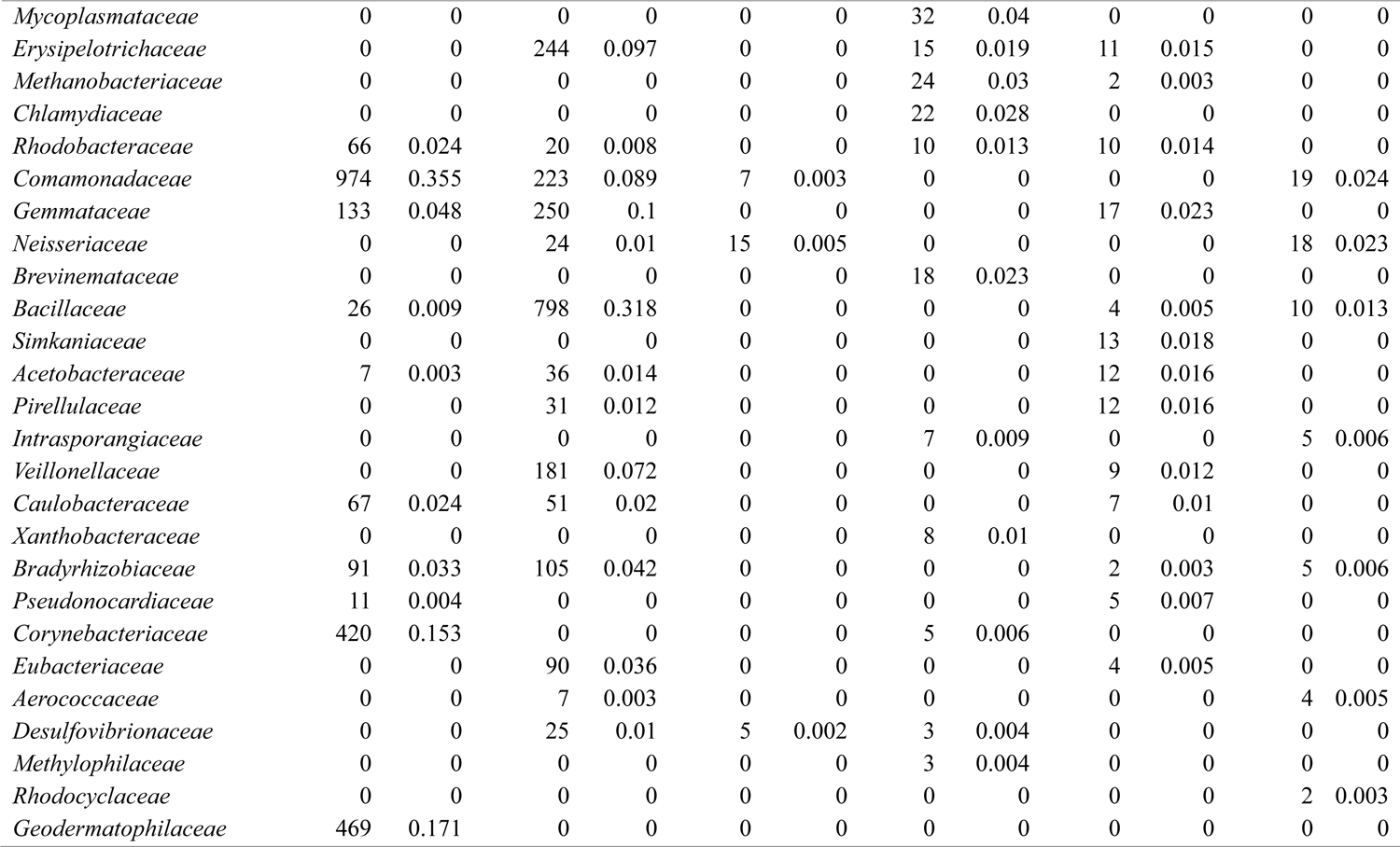
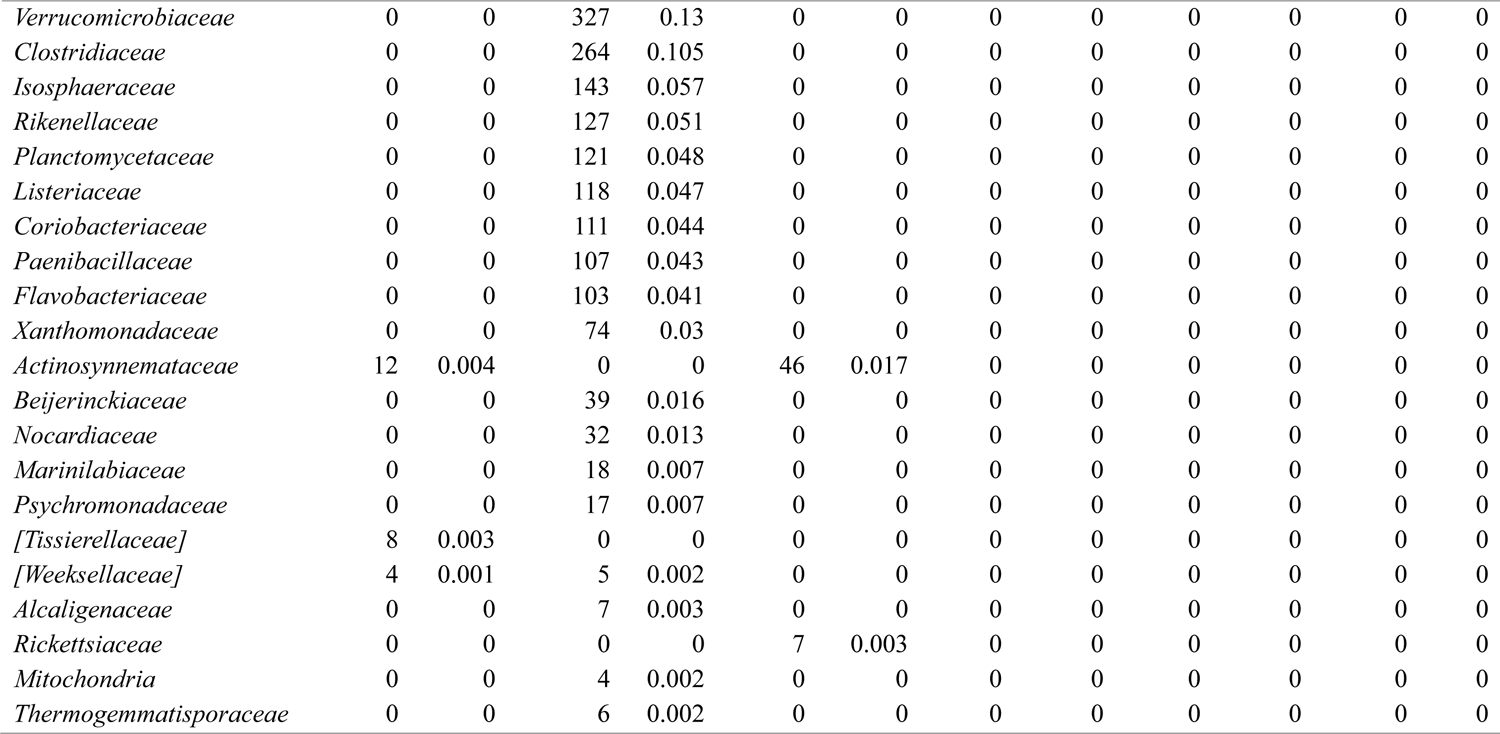
Frequency and relative abundance of each family found in each sample of both healthy and diseased *T. tamrboides,* whereby H1, H2 and H3 stands for healthy *T. tambroides* while D1, D2 and D3 stands for diseased *T. tambroides*.

**Table S6.**
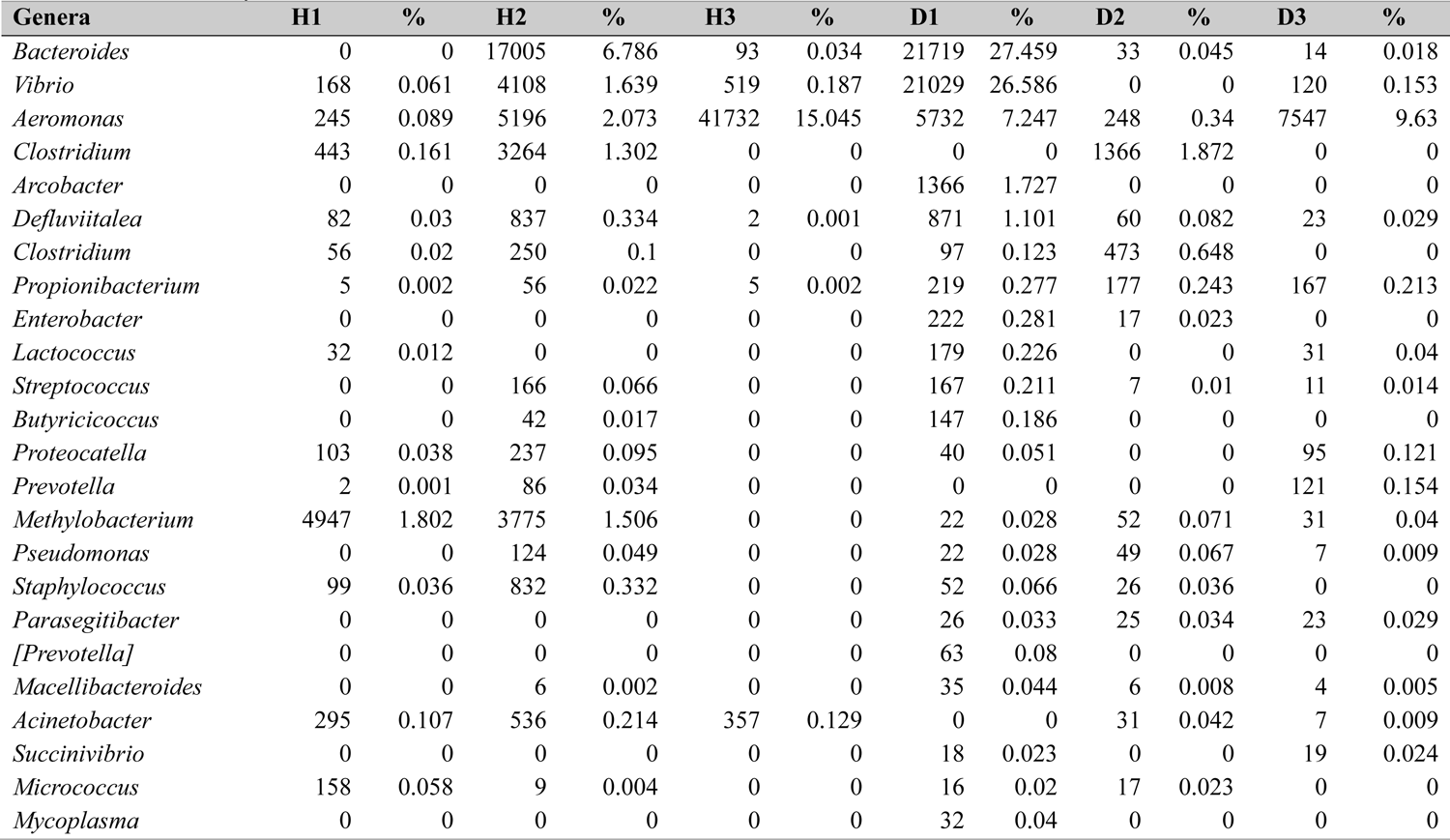
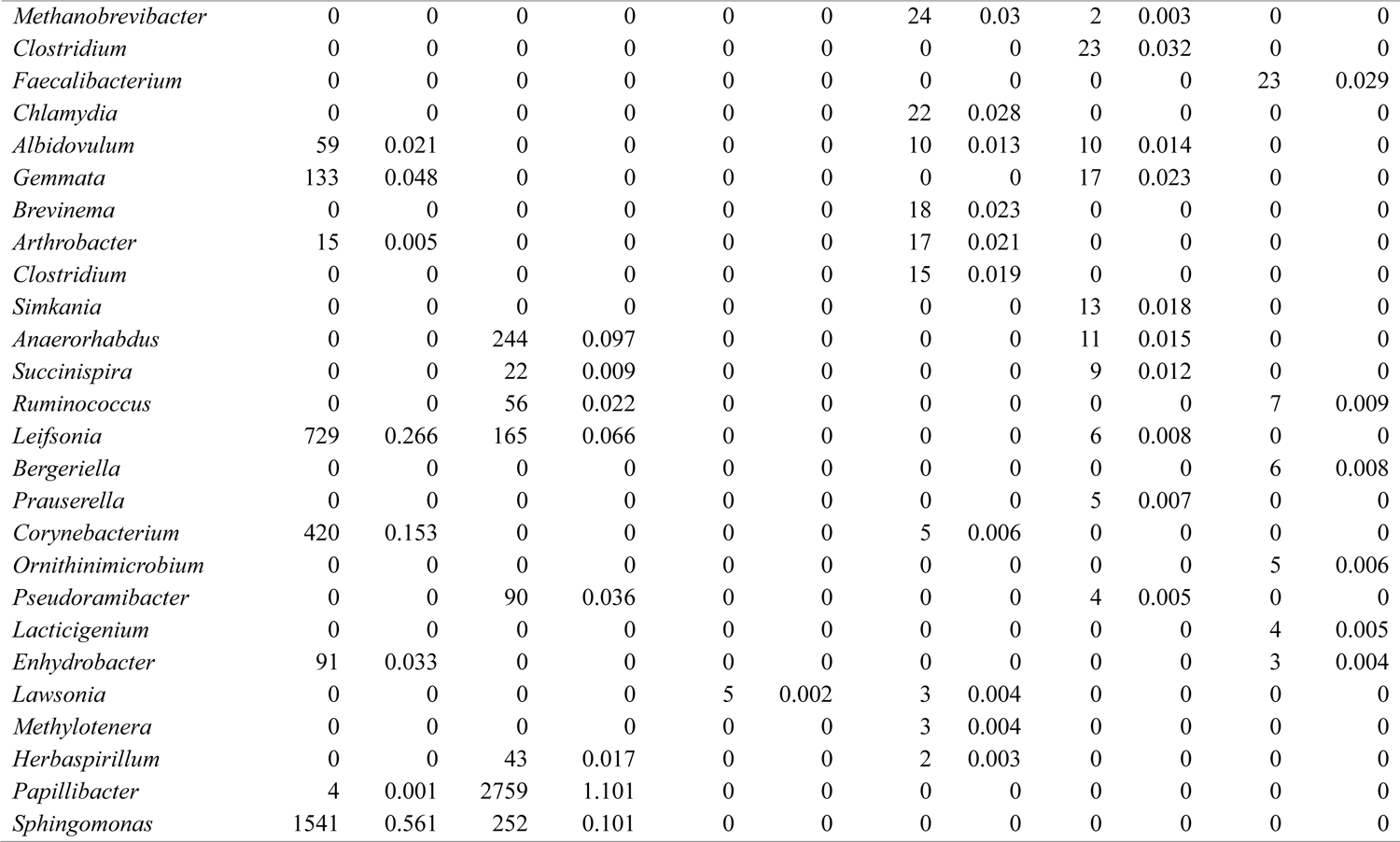
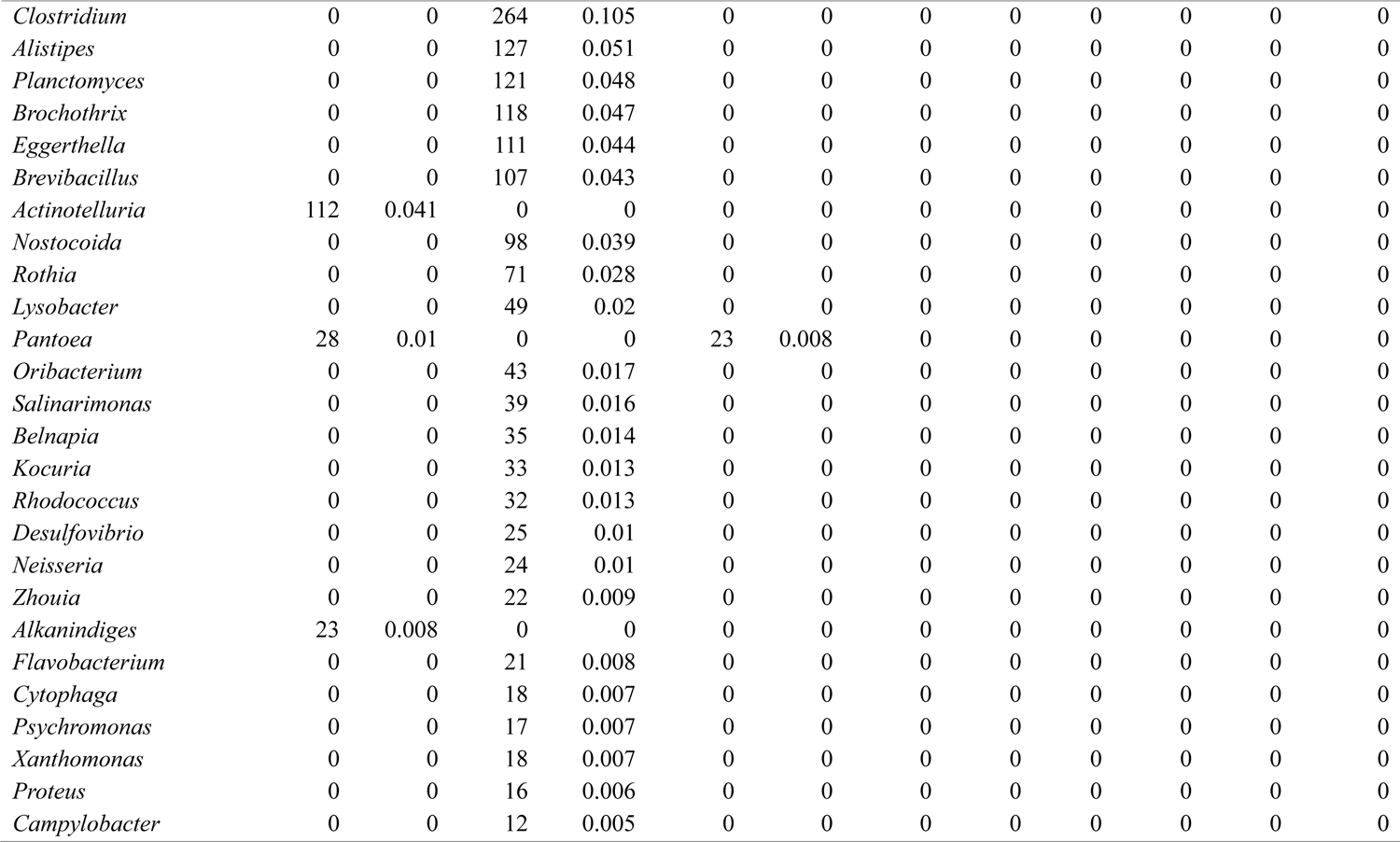
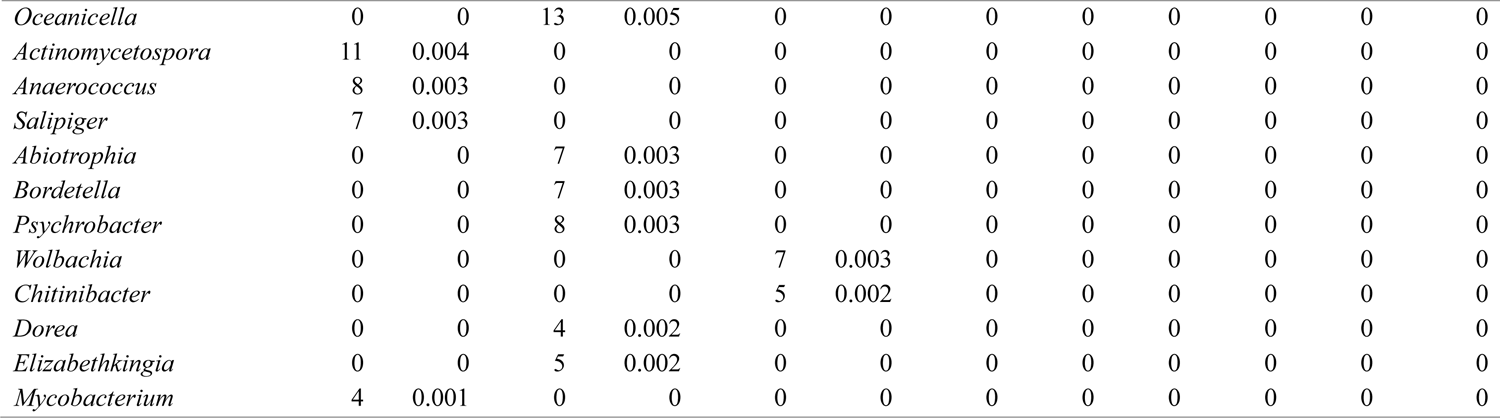
Frequency and relative abundance of each genera found in each sample of both healthy and diseased *T. tamrboides,* whereby H1, H2 and H3 stands for healthy *T. tambroides* while D1, D2 and D3 stands for diseased *T. tambroides*.

**Table S7.**
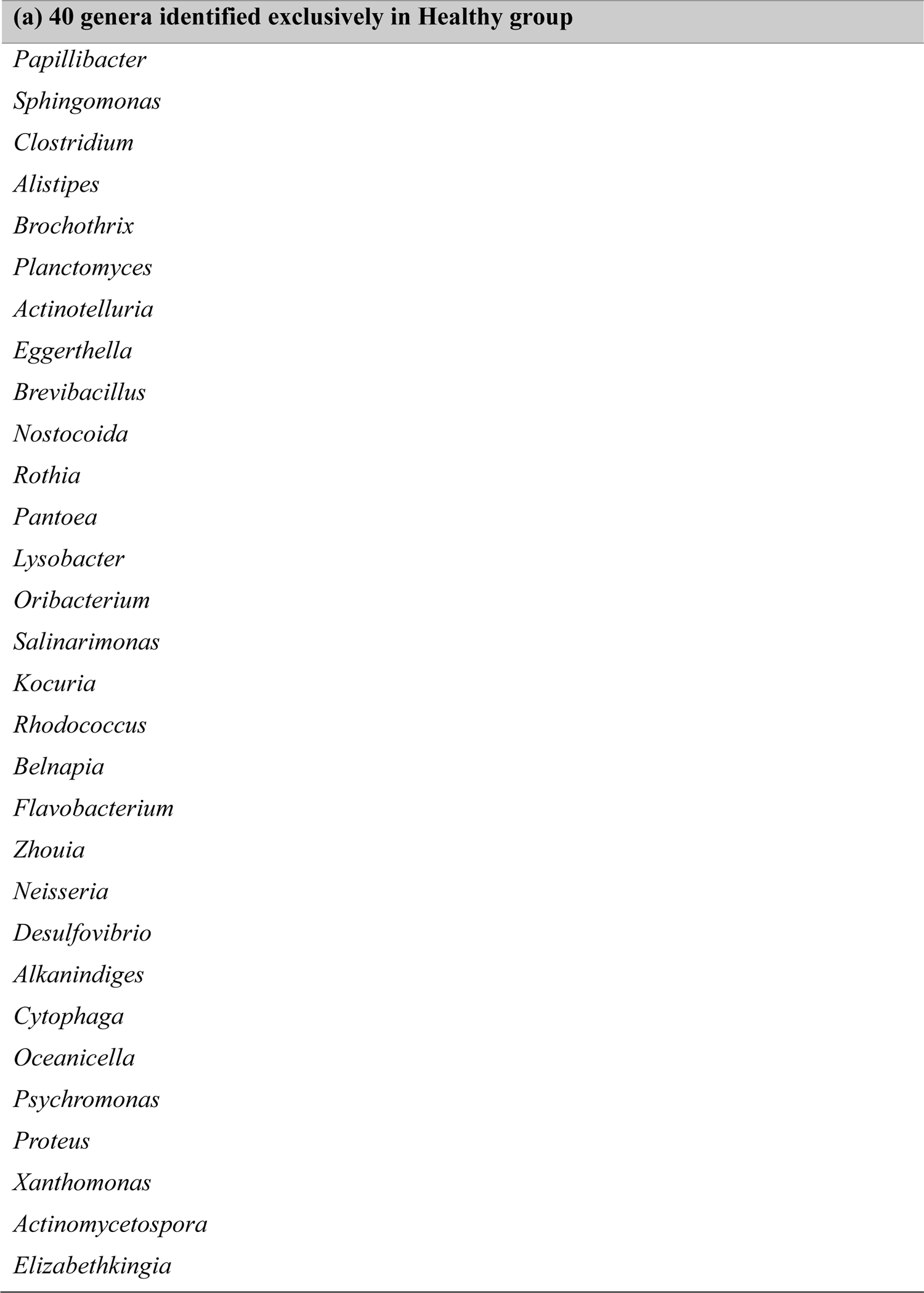
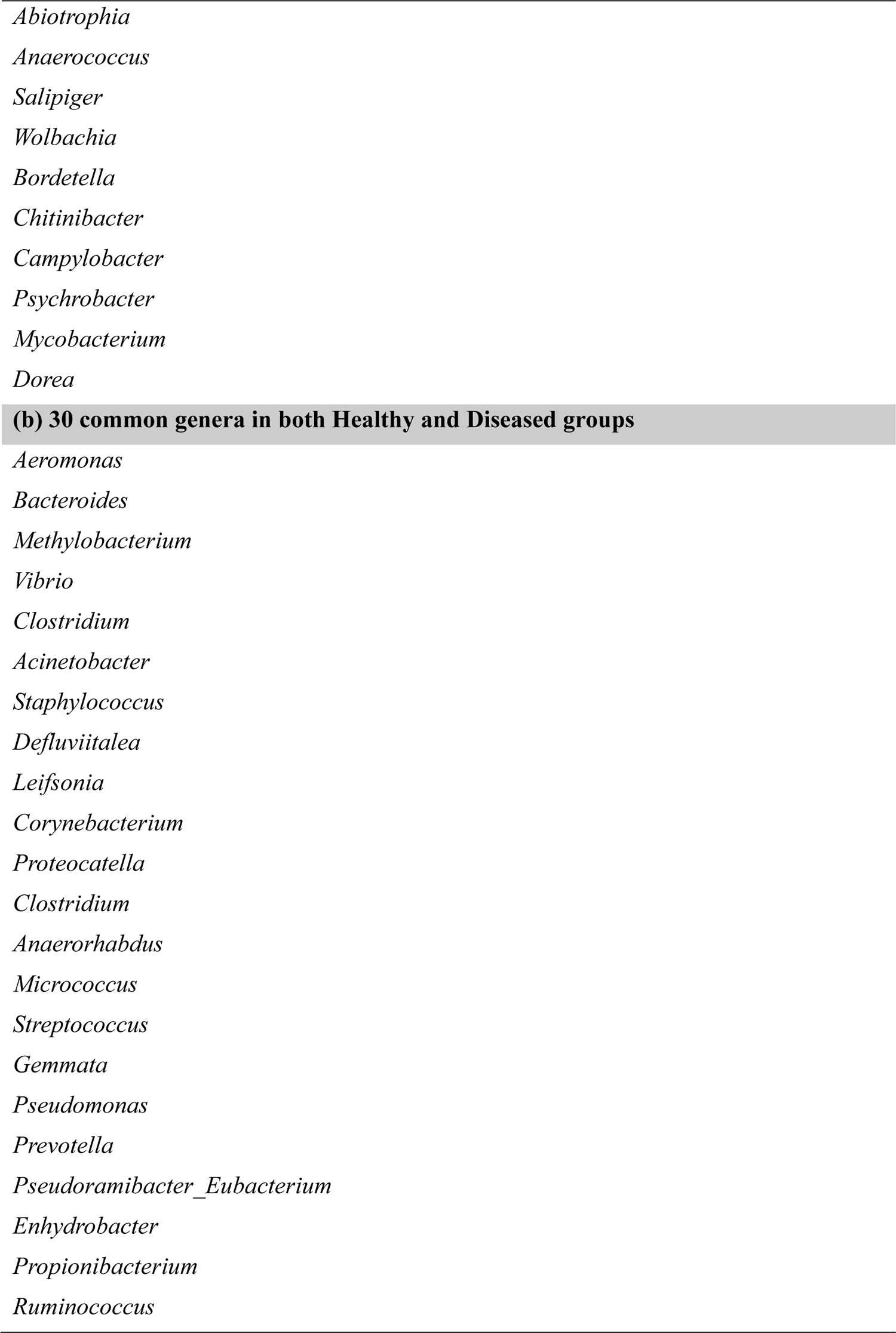
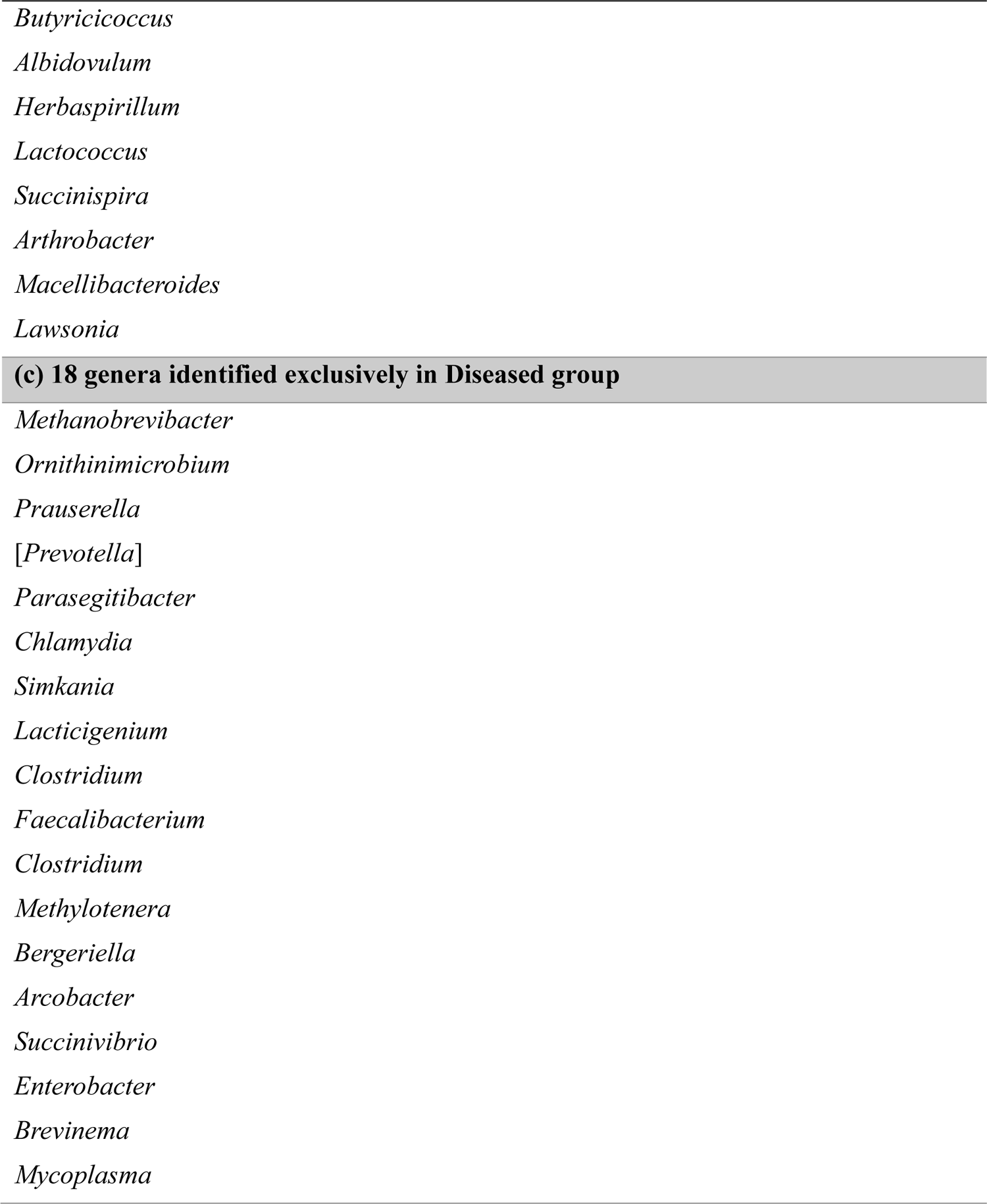
List of genera identified exclusively: Table S4(a) listed on 40 genera unique to healthy gut of *T. tambroides*; Table S4(b) listed on 30 common genera found in both healthy and diseased gut of *T. tambroides*; Table S4(c) listed on 18 genera unique to diseased gut of *T. tambroides*.

**Table S8.**
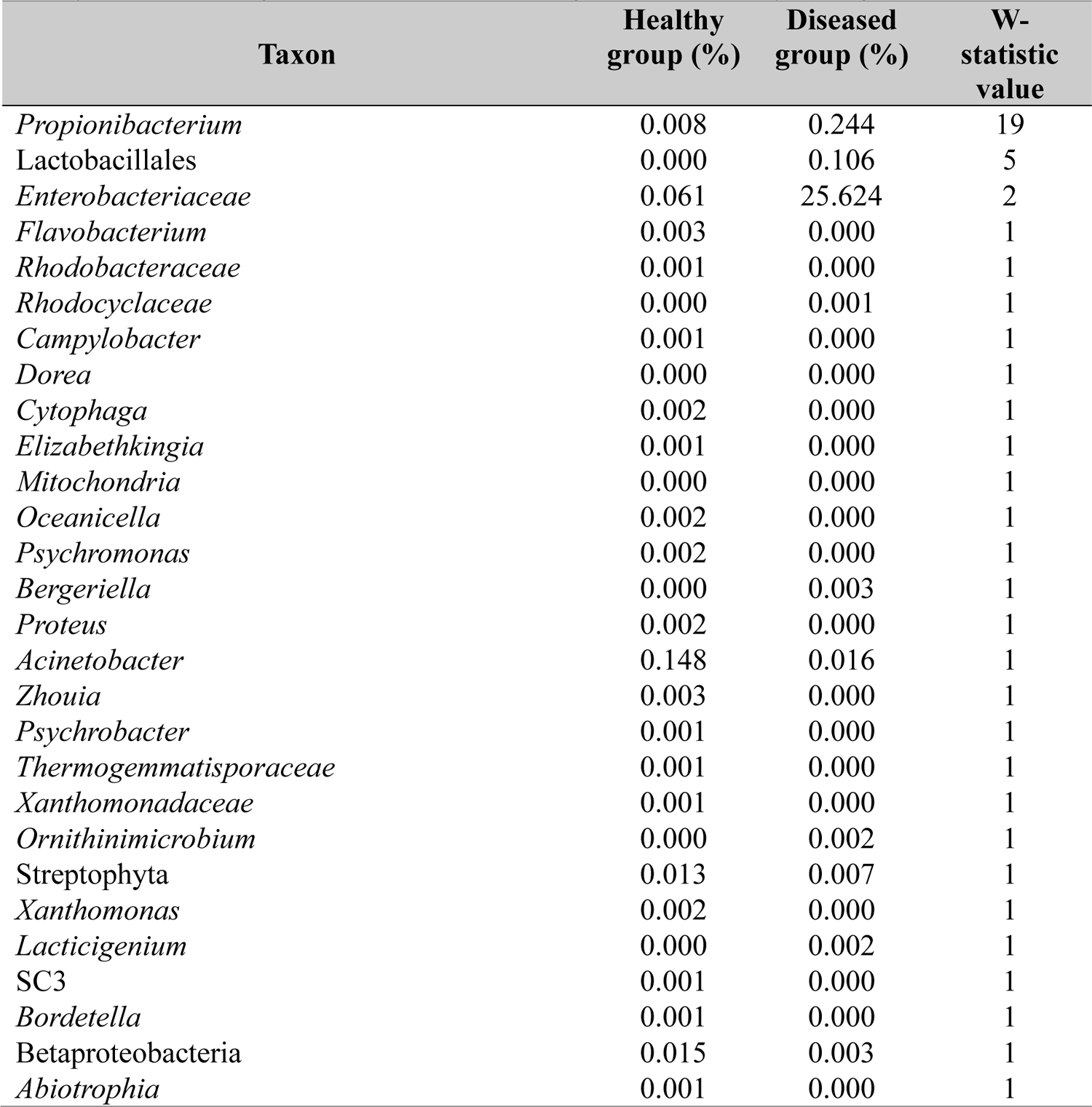
List of observed bacteria that was found to be significant different among both healthy and diseased gut of *T. tambroides* through ANCOM analysis at genus level.

## REFERENCES

1. Açik, M. N., Yüksel, H., Ulucan, A., and Çetinkaya, B. (2016). The first experimental research on the pathogenicity of *Arcobacter butzleri* in zebrafish. Veterinary Microbiology, 189, 32–38.

2. Aly, S., and Ismail, M. (2016). Characteristics of infectious dropsy from an epizootic of cultured common carp (*Cyprinus carpio* L.) with special investigation to swim bladder lesions. Suez Canal Veterinary Medical Journal, 21(1), 185–195.

3. Austin, B. and Austin, D. A. (2012). Bacterial fish pathogens. Springer, The Netherlands. pp. 482.

4. Bolyen, E., Rideout, J. R., Dillon, M. R., Bokulich, N. A., Abnet, C. C., Al-Ghalith, G. A. … and Caporaso, J. G. (2019). Reproducible, interactive, scalable and extensible microbiome data science using QIIME 2. Nature Biotechnology, 37(8), 852–857.

5. Buffie, C. G., Jarchum, I., Equinda, M., Lipuma, L., Gobourne, A., Viale, A. … and Pamer, E. G. (2012). Profound alterations of intestinal microbiota following a single dose ofclindamycin results in sustained susceptibility to *Clostridium difficile*-induced colitis. Infection and Immunity, 80(1), 62–73.

6. Burtseva, O., Kublanovskaya, A., Fedorenko, T., Lobakova, E. and Chekanov, K. (2021). Gut microbiome of the White Sea fish revealed by 16S rRNA metabarcoding. Aquaculture, 533, 736175.

7. Butt, R. L. and Volkoff, H. (2019). Gut microbiota and energy homeostasis in fish. Frontiers in Endocrinology, 10, 9.

8. Callahan, B. J., McMurdie, P. J. and Holmes, S. P. (2017). Exact sequence variants should replace operational taxonomic units in marker-gene data analysis. The ISME Journal, 11(12), 2639–2643.

9. Chao, A. (1984). Nonparametric estimation of the number of classes in a population. Scandinavian Journal of Statistics, 265-270.

10. Chiew, I., Salter, A. M. and Lim, Y. S. (2019). The significance of major viral and bacterial diseases in Malaysian aquaculture industry. Pertanika Journal of Tropical Agricultural Science, 42(3).

11. Chin, A. C. and Parkos, C. A. (2006). Neutrophil transepithelial migration and epithelial barrier function in IBD: Potential targets for inhibiting neutrophil trafficking. Annals of the New York Academy of Sciences, 1072(1), 276–287.

12. Chung, H. H. (2018). Real-time polymerase chain reaction (RT-PCR) for the authentication of raw meats. International Food Research Journal, 25(2), 632–638.

13. Corona-Cervantes, K., García-González, I., Villalobos-Flores, L. E., Hernández-Quiroz, F., Piña-Escobedo, A., Hoyo-Vadillo, C. … and García-Mena, J. (2020). Human milk microbiota associated with early colonization of the neonatal gut in Mexican newborns. PeerJ, 8, e9205.

14. Dash, S., Swain, P., Swain, M. M., Nayak, S. K., Behura, A., Nanda, P. K. and Mishra, B. K. (2008). Investigation on infectious dropsy of Indian major carps. Asian Fisheries Science, 21(4), 377–384.

15. D’Auria, G., Peris-Bondia, F., Džunková, M., Mira, A., Collado, M. C., Latorre, A. and Moya, A. (2013). Active and secreted IgA-coated bacterial fractions from the human gut reveal an under-represented microbiota core. Scientific Reports, 3(1), 1–9.

16. DeSantis, T. Z., Hugenholtz, P., Larsen, N., Rojas, M., Brodie, E. L., Keller, K. … and Andersen, G. L. (2006). Greengenes, a chimera-checked 16S rRNA gene database and workbench compatible with ARB. Applied and Environmental Microbiology, 72(7), 5069–5072.

17. DOF. (2017). Annual Fisheries Statistics 2017. Kuala Lumpur: Department of Fisheries Malaysia.

18. DOF. (2018). Annual Fisheries Statistics 2018. Kuala Lumpur: Department of Fisheries Malaysia.

19. Dong, H. T., Nguyen, V. V., Le, H. D., Sangsuriya, P., Jitrakorn, S., Saksmerprome, V. … and Rodkhum, C. (2015). Naturally concurrent infections of bacterial and viral pathogens in disease outbreaks in cultured Nile tilapia (*Oreochromis niloticus*) farms. Aquaculture, 448, 427–435.

20. Dong, H. T., Techatanakitarnan, C., Jindakittikul, P., Thaiprayoon, A., Taengphu, S., Charoensapsri, W. … and Senapin, S. (2017). *Aeromonas jandaei* and *Aeromonas veronii* caused disease and mortality in Nile tilapia, *Oreochromis niloticus* (L.). Journal of Fish Diseases, 40(10), 1395–1403.

21. Egerton, S., Culloty, S., Whooley, J., Stanton, C. and Ross, R. P. (2018). The gut microbiota of marine fish. Frontiers in Microbiology, 9, 873.

22. Elinav, E., Strowig, T., Kau, A. L., Henao-Mejia, J., Thaiss, C. A., Booth, C. J. … and Flavell, R. A. (2011). NLRP6 inflammasome regulates colonic microbial ecology and risk forcolitis. Cell, 145(5), 745–757.

23. Esa, Y., Siraj, S.S., Daud, S.K., Rahim, A., Adha, K., Abdullah, M. T. … and Tan, S. G. (2006). Mitochondrial DNA Diversity of *Tor douronensis* Valenciennes (*Cyprinidae*) in Malaysian Borneo. Pertanika Journal of Tropical Agricultural Science, 29 (1 & 2), 47–55.

24. Fargione, J. E. and Tilman, D. (2005). Diversity decreases invasion via both sampling and complementarity effects. Ecology Letters, 8(6), 604–611.

25. García-López, R., Cornejo-Granados, F., Lopez-Zavala, A. A., Sánchez-López, F., Cota Huízar, A., Sotelo-Mundo, R. R. … and Ochoa-Leyva, A. (2020). Doing more with less: A comparison of 16S hypervariable regions in search of defining the shrimp microbiota. Microorganisms, 8(1), 134.

26. Ghanbari, M., Jami, M., Shahraki, H. and Domig, K. J. (2019). Characterization of the gut microbiota in the Anjak (Schizocypris altidorsalis), a native fish species of Iran. BioRxiv, 735241.

27. Hagi, T. and Hoshino, T. (2009). Screening and characterization of potential probiotic lactic acid bacteria from cultured common carp intestine. Bioscience, Biotechnology, and Biochemistry, 73(7), 1479–1483.

28. Hamid, R., Ahmad, A. and Usup, G. (2016). Pathogenicity of *Aeromonas hydrophila* isolated from the Malaysian Sea against coral (*Turbinaria* sp.) and sea bass (Lates calcarifer). Environmental Science and Pollution Research, 23(17), 17269–17276.

29. Ho, H. T., Lipman, L. J. and Gaastra, W. (2006). *Arcobacter*, what is known and unknown about a potential foodborne zoonotic agent!. Veterinary Microbiology, 115(1-3), 1–13.

30. Hovda, M. B., Lunestad, B. T., Fontanillas, R. and Rosnes, J. T. (2007). Molecular characterisation of the intestinal microbiota of farmed Atlantic salmon (*Salmo salar* L.). Aquaculture, 272(1-4), 581–588.

31. Ingram, B., Sungan, S., Gooley, G., Sim, S. Y., Tinggi, D. and De Silva, S. S. (2005). Induced spawning, larval development and rearing of two indigenous Malaysian mahseer, Tor tambroides and T. douronensis. Aquaculture Research, 36(10), 983–995.

32. Ishak, S. D., Kamarudin, M. S., Ramezani-Fard, E., Saad, C. R. and Yusof, Y. A. (2016). Effects of varying dietary carbohydrate levels on growth performance, body composition and liver histology of Malaysian mahseer fingerlings (*Tor tambroides*). Journal of Environmental Biology, 37, 755–764.

33. Kamada, N., Seo, S. U., Chen, G. Y. and Núñez, G. (2013). Role of the gut microbiota in immunity and inflammatory disease. Nature Reviews Immunology, 13(5), 321–335.

34. Kamarudin, M. S. (2015). Feeding and nutritional requirements of young fish. Inaugural Lecture Series. Selangor: Universiti Putra Malaysia.

35. Kamarudin, M. S., Ramezani-Fard, E., Saad, C. R. and Harmin, S. A. (2012). Effects of dietary fish oil replacement by various vegetable oils on growth performance, body composition and fatty acid profile of juvenile Malaysian mahseer, *Tor tambroides*. Aquaculture Nutrition, 18(5), 532–543.

36. Karadas, G., Sharbati, S., Hänel, I., Messelhäußer, U., Glocker, E., Alter, T. and Gölz, G. (2013). Presence of virulence genes, adhesion and invasion of *Arcobacter butzleri*. Journal of Applied Microbiology, 115(2), 583–590.

37. Kottelat, M., Pinder, A. and Harrison, A. (2018). Tor tambroides. The IUCN Red List of Threatened Species 2018.

38. Lane, D. J. (1991). Nucleic acid sequencing techniques in bacterial systematics.

39. Lau, M. M. L., Lim, L. W. K., Ishak, S. D., Abol-Munafi, A. B. and Chung, H. H. (2021). A review on the emerging asian aquaculture fish, the Malaysian Mahseer (*Tor tambroides*): Current status and the way forward. In Proceedings of the Zoological Society. pp. 1–11.

40. Li, T., Li, H., Gatesoupe, F. J., She, R., Lin, Q., Yan, X. … and Li, X. (2017). Bacterial signatures of “Red-Operculum” disease in the gut of crucian carp (*Carassius auratus*). Microbial Ecology, 74(3), 510–521.

41. Li, T., Long, M., Ji, C., Shen, Z., Gatesoupe, F. J., Zhang, X. … and Li, A. (2016). Alterations of the gut microbiome of largemouth bronze gudgeon (*Coreius guichenoti*) suffering from furunculosis. Scientific Reports, 6(1), 1–9.

42. Lim L. W. K., Chung, H. H., Lau, M. M. L., Aziz, F. and Gan, H. M. (2021). Improving the phylogenetic resolution of Malaysian and Javan mahseer (*Cyprinidae*), *Tor tambroides* and *Tor tambra*: Whole mitogenomes sequencing, phylogeny and potentialmitogenome markers. Gene, 791, 145708.

43. Liu, H., Guo, X., Gooneratne, R., Lai, R., Zeng, C., Zhan, F. and Wang, W. (2016). The gut microbiome and degradation enzyme activity of wild freshwater fishes influenced by their trophic levels. Scientific Reports, 6(1), 1–12.

44. Long, X., Deng, S., Mattner, J., Zang, Z., Zhou, D., McNary, N. … and Savage, P. B. (2007). Synthesis and evaluation of stimulatory properties of *Sphingomonadaceae* glycolipids. Nature Chemical Biology, 3(9), 559–564.

45. Lopamudra, M.B. and Nayak, S.K. (2020). Characterization and serosurveillance of *Aeromonas hydrophila* infection in disease affected freshwater fishes. Journal of Fisheries and Aquatic Science, 5, 39–44.

46. Lozupone, C. A., Hamady, M., Kelley, S. T. and Knight, R. (2007). Quantitative and qualitative β diversity measures lead to different insights into factors that structure microbial communities. Applied and Environmental Microbiology, 73(5), 1576–1585.

47. Mandal, S., Van Treuren, W., White, R. A., Eggesbø, M., Knight, R. and Peddada, S. D. (2015). Analysis of composition of microbiomes: A novel method for studying microbial composition. Microbial Ecology in Health and Disease, 26(1), 27663.

48. Mathias, J. R., Perrin, B. J., Liu, T. X., Kanki, J., Look, A. T. and Huttenlocher, A. (2006). Resolution of inflammation by retrograde chemotaxis of neutrophils in transgenic zebrafish. Journal of Lleukocyte Bbiology, 80(6), 1281–1288.

49. Misieng, J. D., Kamarudin, M. S. and Musa, M. (2011). Optimum dietary protein requirement of Malaysian mahseer *(Tor tambroides*) fingerling. Pakistan Journal of Biological Sciences, 14(3), 232–235.

50. Namba, A., Mano, N. and Hirose, H. (2007). Phylogenetic analysis of intestinal bacteria and their adhesive capability in relation to the intestinal mucus of carp. Journal of Applied Microbiology, 102(5), 1307–1317.

51. Narrowe, A. B., Albuthi-Lantz, M., Smith, E. P., Bower, K. J., Roane, T. M., Vajda, A. M. and Miller, C. S. (2015). Perturbation and restoration of the fathead minnow gut microbiome after low-level triclosan exposure. Microbiome, 3(1), 1–18.

52. Nayak, S. K. (2010). Role of gastrointestinal microbiota in fish. Aquaculture Research, 41(11), 1553–1573.

53. Nearing, J. T., Douglas, G. M., Comeau, A. M. and Langille, M. G. (2018). Denoising the denoisers: An independent evaluation of microbiome sequence error-correction approaches. PeerJ, 6, e5364.

54. Frank, D. N., Amand, A. L. S., Feldman, R. A., Boedeker, E. C., Harpaz, N. and Pace, N. R. (2007). Molecular-phylogenetic characterization of microbial community imbalances in human inflammatory bowel diseases. Proceedings of the National Academy of Sciences, 104(34), 13780–13785.

55. Ng, W. K., Abdullah, N. and De Silva, S. S. (2008). The dietary protein requirement of the Malaysian mahseer, *Tor tambroides* (Bleeker), and the lack of protein-sparing action by dietary lipid. Aquaculture, 284(1-4), 201–206.

56. Nie, L., Zhou, Q. J., Qiao, Y. and Chen, J. (2017). Interplay between the gut microbiota and immune responses of ayu (*Plecoglossus altivelis*) during *Vibrio anguillarum* infection. Fish & Shellfish Immunology, 68, 479–487.

57. O’Hara, A. M. and Shanahan, F. (2006). The gut flora as a forgotten organ. EMBO Reports, 7(7), 688–693.

58. Petty, B. D., Francis-Floyd, R. and Yanong, R. P. (2012). Spring viremia of carp. EDIS, 2012(9).

59. Pinder, A. C., Britton, J. R., Harrison, A. J., Nautiyal, P., Bower, S. D., Cooke, S. J. … and Raghavan, R. (2019). Mahseer (*Tor* spp.) fishes of the world: Status, challenges and opportunities for conservation. Reviews in Fish Biology and Fisheries, 29, 417–452.

60. Prodan, A., Tremaroli, V., Brolin, H., Zwinderman, A. H., Nieuwdorp, M. and Levin, E. (2020). Comparing bioinformatic pipelines for microbial 16S rRNA amplicon sequencing. PLoS One, 15(1), e0227434.

61. Ramezani-Fard, E., Kamarudin, M. S., Harmin, S. A. and Saad, C. R. (2012). Dietary saturated and omega-3 fatty acids affect growth and fatty acid profiles of Malaysian Mahseer. European Journal of Lipid Science and Technology, 114(2), 185–193.

62. Shannon, C. E. (2001). A mathematical theory of communication. ACM SIGMOBILE Mobile Computing and Communications Review, 5(1), 3–55.

63. She, R., Li, T. T., Luo, D., Li, J. B., Yin, L. Y., Li, H. … and Yan, Q. G. (2017). Changes in the intestinal microbiota of gibel carp (*Carassius gibelio*) associated with cyprinid herpesvirus 2 (CyHV-2) infection. Current Microbiology, 74(10), 1130–1136.

64. Simpson, E. H. (1949). Measurement of diversity. Nature, 163(4148), 688–688.

65. Siraj, S.S., Esa, Y.B., Keong, B.P. and Daud, S.K. (2007). Genetic characterization of the two colour-type of Kelah. Malaysian Applied Biology, 36, 23–29.

66. Skrodenytė-Arbačiauskienė, V. (2000). Proteolytic activity of the roach (*Rutilus rutilus* L.) intestinal microflora. Acta Zoologica Lituanica, 10(3), 69–77.

67. Stecher, B. (2015). The roles of inflammation, nutrient availability and the commensal microbiota in enteric pathogen infection. Microbiology Spectrum, 3(3), 3–3.

68. Tan, C. K., Natrah, I., Suyub, I. B., Edward, M. J., Kaman, N. and Samsudin, A. A. (2019). Comparative study of gut microbiota in wild and captive Malaysian Mahseer (*Tor tambroides*). Microbiologyopen, 8(5), e00734.

69. Tarnecki, A. M., Burgos, F. A., Ray, C. L. and Arias, C. R. (2017). Fish intestinal microbiome: Diversity and symbiosis unravelled by metagenomics. Journal of Applied Microbiology, 123(1), 2–17.

70. Tran, N. T., Zhang, J., Xiong, F., Wang, G. T., Li, W. X. and Wu, S. G. (2018). Altered gut microbiota associated with intestinal disease in grass carp (*Ctenopharyngodon idellus*). World Journal of Microbiology and Biotechnology, 34(6), 1–9.

71. Vatsos, I. N. (2017). Standardizing the microbiota of fish used in research. Laboratory Animals, 51(4), 353–364.

72. Wallace, J. L., Syer, S., Denou, E., de Palma, G., Vong, L., McKnight, W. … and Ongini, E. (2011). Proton pump inhibitors exacerbate NSAID-induced small intestinal injury by inducing dysbiosis. Gastroenterology, 141(4), 1314–1322.

73. Walton, S. E., Gan, H. M., Raghavan, R., Pinder, A. C. and Ahmad, A. (2017). Disentangling the taxonomy of the mahseers (*Tor* spp.) of Malaysia: An integrated approach using morphology, genetics and historical records. Reviews in Fisheries Science and Aquaculture, 25, 171–183.

74. Wu, S., Wang, G., Angert, E. R., Wang, W., Li, W. and Zou, H. (2012). Composition, diversity, and origin of the bacterial community in grass carp intestine. PloS one, 7(2), e30440.

75. Xiong, J. B., Nie, L. and Chen, J. (2019). Current understanding on the roles of gut microbiota in fish disease and immunity. Zoological Research, 40(2), 70.

76. Zeng, M. Y., Inohara, N. and Nuñez, G. (2017). Mechanisms of inflammation-driven bacterial dysbiosis in the gut. Mucosal Immunology, 10(1), 18–26.

